# Sexual differences in genetic architecture in UK Biobank

**DOI:** 10.1101/2020.07.20.211813

**Authors:** Elena Bernabeu, Oriol Canela-Xandri, Konrad Rawlik, Andrea Talenti, James Prendergast, Albert Tenesa

**Affiliations:** The Roslin Institute, Royal (Dick) School of Veterinary Studies, The University of Edinburgh, Easter Bush Campus, Midlothian, EH25 9RG, UK; MRC Human Genetics Unit at the MRC Institute of Genetics and Molecular Medicine, University of Edinburgh, Western General Hospital, Crewe Road South, Edinburgh, EH4 2XU, UK

## Abstract

Sex is arguably the most important differentiating characteristic in most mammalian species, separating populations into different groups, with varying behaviors, morphologies, and physiologies based on their complement of sex chromosomes. In humans, despite males and females sharing nearly identical genomes, there are differences between the sexes in complex traits and in the risk of a wide array of diseases. Gene by sex interactions (GxS) are thought to account for some of this sexual dimorphism. However, the extent and basis of these interactions are poorly understood.

Here we provide insights into both the scope and mechanism of GxS across the genome of circa 450,000 individuals of European ancestry and 530 complex traits in the UK Biobank. We found small yet widespread differences in genetic architecture across traits through the calculation of sex-specific heritability, genetic correlations, and sex-stratified genome-wide association studies (GWAS). We also found that, in some cases, sex-agnostic GWAS efforts might be missing loci of interest, and looked into possible improvements in the prediction of high-level phenotypes. Finally, we studied the potential functional role of the dimorphism observed through sex-biased eQTL and gene-level analyses.

This study marks a broad examination of the genetics of sexual dimorphism. Our findings parallel previous reports, suggesting the presence of sexual genetic heterogeneity across complex traits of generally modest magnitude. Our results suggest the need to consider sex-stratified analyses for future studies in order to shed light into possible sex-specific molecular mechanisms.

## INTRODUCTION

In recent years, there has been growing evidence of common genetic variation having different effects on males and females^1,2^. This, along with sex-biases observed in the human transcriptome^3–10^, the presence of a distinct hormone milieu in each sex, and differential environmental pressures arising from gender societal roles^1,11^, has led to an increased study of the potential importance of GxS interactions to understand the underlying biology of complex traits, including the estimation of disease risk. Previous studies have investigated differences in heritability between the sexes (h^2^)^12–14^, departure of genetic correlations from 1 (r_g_)^12,14–18^, and performed sex-stratified genome-wide association studies (GWAS) to directly assess differences in the effects of genetic variants between the sexes^6,19–26^. These studies have however been limited with regards to the number of traits studied or statistical power. Furthermore, insights into how differences in genetic architecture translate into differences in complex traits have been lacking. Accounting for sexual dimorphism is of great importance as sex-agnostic analyses could potentially be masking sex-specific effects, and which could - if better understood - lead to better personalized treatment and an improved understanding of the biological mechanisms driving sexual dimorphism^16,27^.

The objective of the current study was to assess both the existence and scope of GxS interactions in the human genome by estimating sex-specific heritability and genetic correlations, as well as performing sex-stratified GWAS analyses. To this end, we analysed 530 traits using 450K individuals of European ancestry from the UK Biobank. Furthermore, we evaluated the potential of improving trait predictions using sex-stratified polygenic scores (PGS), as well as looked into the possibility of missing loci of interest in non-sex stratified studies. Finally, to shed light on the downstream effects of sex-differences in genetic architecture, we performed a functional *in silico* analysis.

## RESULTS

### Data overview

Using the July 2017 release of the UK Biobank dataset, we performed sex-stratified GWASs and partition of variance analyses for 530 traits (446 binary and 84 non-binary, Table S1) within 452,264 individuals of European ancestry (245,494 females and 206,770 males) using DISSECT^28^. Linear Mixed Models (LMMs) were fitted for each phenotype by sex. We then tested the association of 9,072,751 autosomal and 17,364 X-chromosome genetic variants, obtaining estimates of the genetic effects of each variant in each sex. In our quality control (QC) stage we excluded genetic variants with a minimum allele frequency (MAF) < 10% in the analysis of binary traits due to the general limited number of cases available, thus reducing the number of variants considered for these traits to 4,229,346 autosomal and 7,227 genotyped X-chromosome genetic variants (see Methods). The results of the autosomal analyses were used to estimate sex-stratified genetic parameters (such as heritability) and genetic correlations. We then tested for differences in these genetic parameters and between the effects of genetic variants estimated within sexes (see Methods).

### Heritability differences between the sexes

Heritability (here referring to SNP heritability^29^) is defined as the fraction of the variation of a trait that can be explained by the additive effects of genetic variation. A difference in heritability between the sexes would entail a difference in the fraction of the variance of a trait that is accounted for by the genotype, and thus a possible difference in the underlying genetic mechanisms of said trait. Out of the 530 traits studied, 41/84 (48.88%) non-binary traits and 30/446 (6.73%) binary traits showed significant differences in their heritability between the sexes (FDR corrected *p*, termed *q*, < 0.05, Figure 1, Table S1). Of these, a total of 25/41 (60.98%) non-binary and 14/30 (46.67%) binary traits had a larger heritability in males than in females.

**Figure 1.**
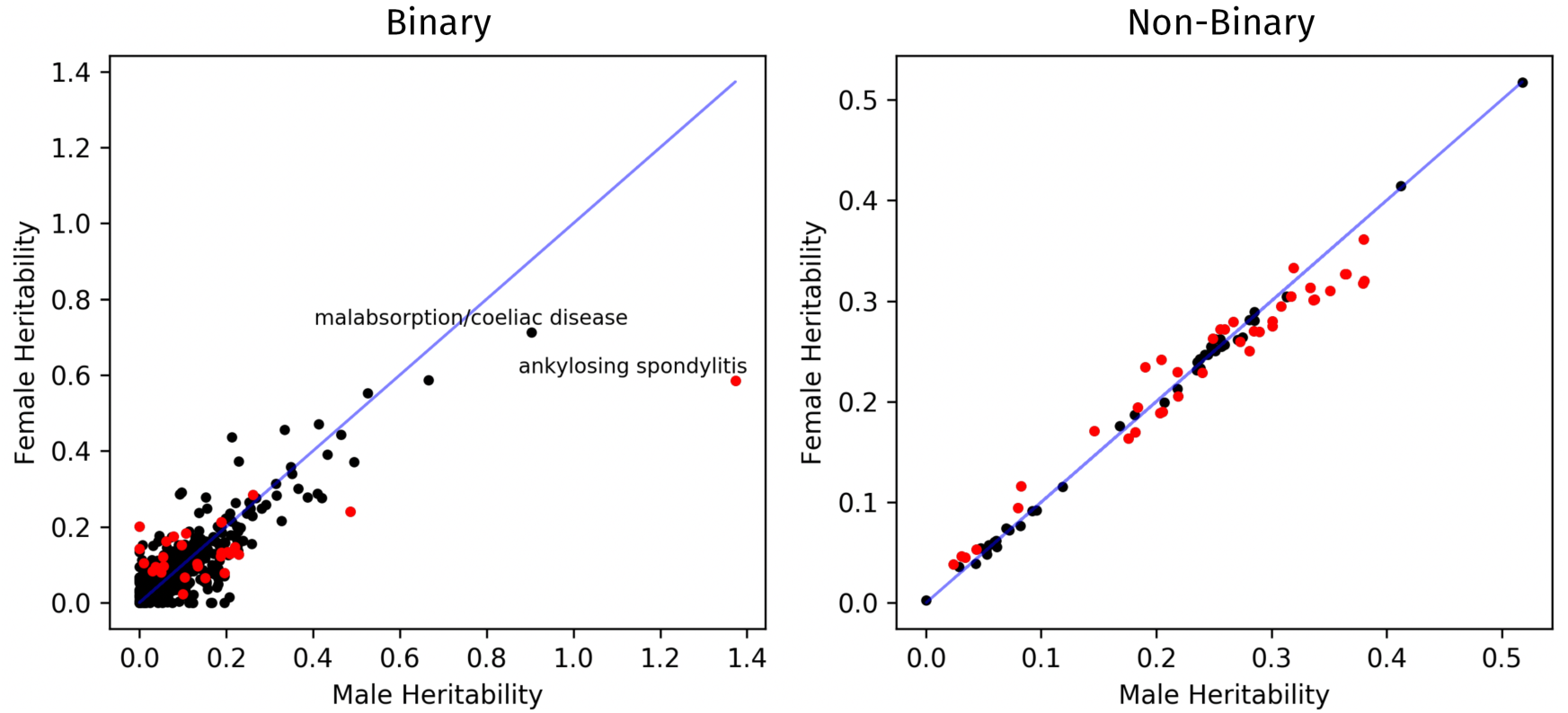
Scatterplot of male heritability estimates against female heritability estimates for binary traits (on the left) and non-binary traits (on the right). Each point represents a trait, which is marked in red when heritability between the sexes is significantly different at a threshold of *q* < 0.05. Note that for the binary traits, heritability estimates were considered on the liability scale, which led to some estimates over 1 (see Methods).

Although differences in the heritability of traits between the sexes can offer potential insights into genetic differences, one must also consider that these could arise due to differences in environmental variances. Hence, we looked for differences in genetic variance between the sexes. We found that 65/84 non-binary (77.38%) and 136/446 binary traits (30.49%) showed significant differences in the amount of genetic variance estimated in each sex (Figure S1, Figure S2A, Table S1).

Finally, we observed significant differences in evolvability (a measure of the ability to undergo adaptation) between sexes for 56/84 non-binary (66.67%) and 35/446 binary (7.85%) (*q* < 0.05, Figure S2B, Table S1). These estimates offer further evidence for differences in the underlying genetic architecture of the traits considered, paralleling previous reports at smaller scales for traits including height, waist-hip circumference ratio and weight^12–14^.

### Genetic correlations indicate the presence of sex by genotype interactions

Genetic correlations (r_g_) between two sub-groups of the population are usually interpreted as a measure of shared underlying genetics, and are a means to estimate the size of putative genotype by group interactions. Genetic correlations between the sexes can thus offer insights into the common genetic control of complex traits and diseases of males and females.

We obtained genetic correlations between the sexes for a total of 83 non-binary and 77 binary traits with over 5,000 cases using LD score regression (LDSC, Table S1)^30^ which met our QC criteria (see Methods). Genetic correlations ranged from 0.716 to 0.996 for non-binary traits and from 0.226 to 1.099 for binary traits (note that with heritability close to zero application of LDSC can result in r_g_ exceeding the theoretically valid range [-1, 1]^31^). A total of 58/83 (69.88%) non-binary traits and 11/77 (14.29%) binary traits had an r_g_ significantly different from 1 (*q* < 0.05, Figure 2A). These included binary traits like hernia (r_g_ = 0.59, *q =* 4.04 x 10^−10^), eczema (r_g_ = 0.61, *q* = 0.04) and gastric reflux (r_g_ = 0.67, *q* = 0.02), and non-binary traits like waist-hip circumference ratio (r_g_ = 0.72, *q* = 8.43 x 10^−37^) and alcohol intake frequency (r_g_ = 0.85, *q* = 7.49 x 10^−9^). Our r_g_ estimates for several non-binary traits were in line with what has previously been published (Table S2, Figure 2B)^12,14,15^.

**Figure 2.**
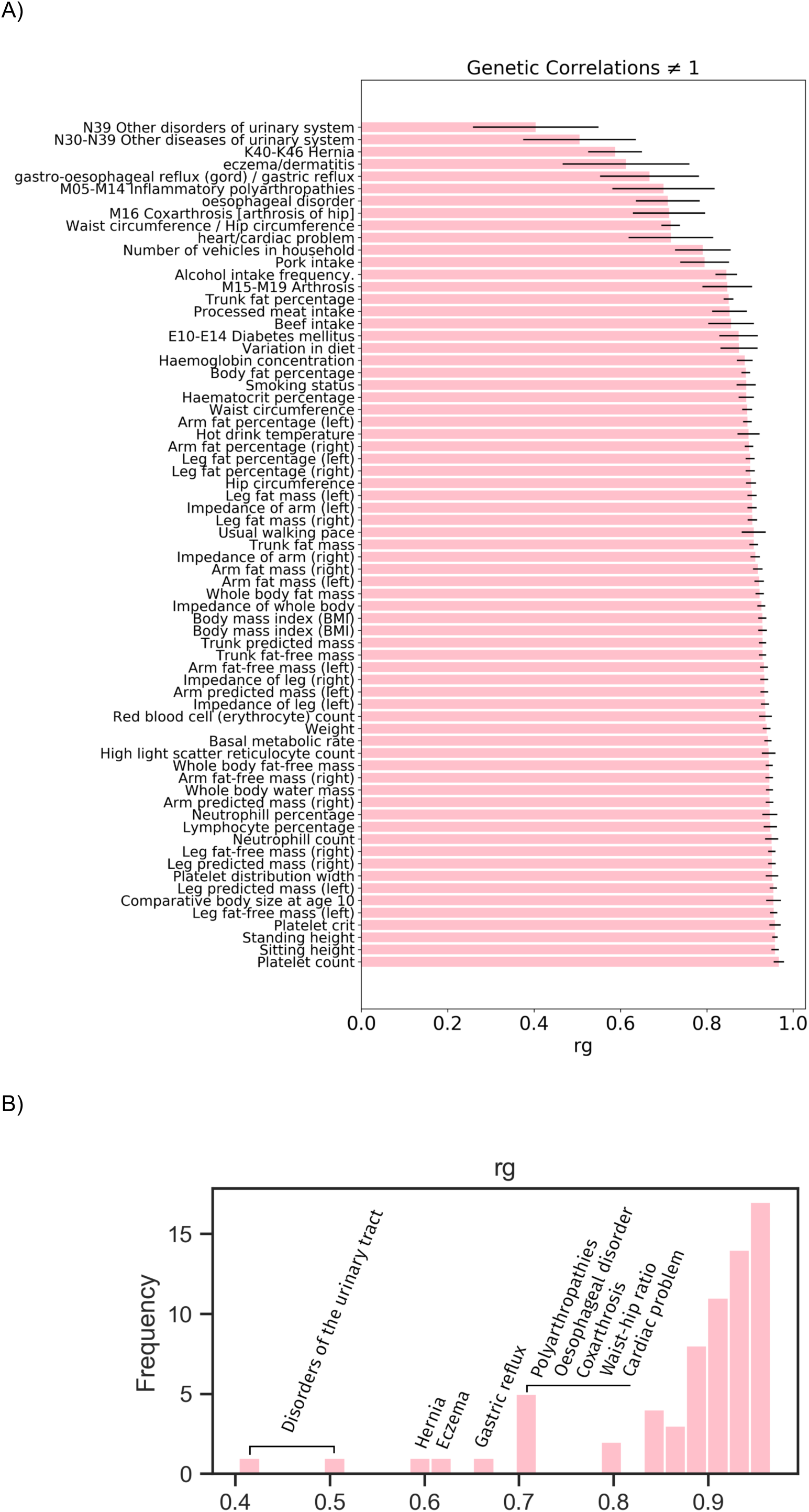
(A) Barplot of genetic correlations (r_g_) between the sexes for traits that were found to have an r_g_ significantly different from one (*q* < 0.05). Black bars indicate the standard errors of the r_g_ estimates. (B) Histogram of genetic correlations that were found to be significantly different from one (*q* < 0.05) for both binary and non-binary traits, which shows a large predominance of r_g_ estimates close to 1.

**Figure 3.**
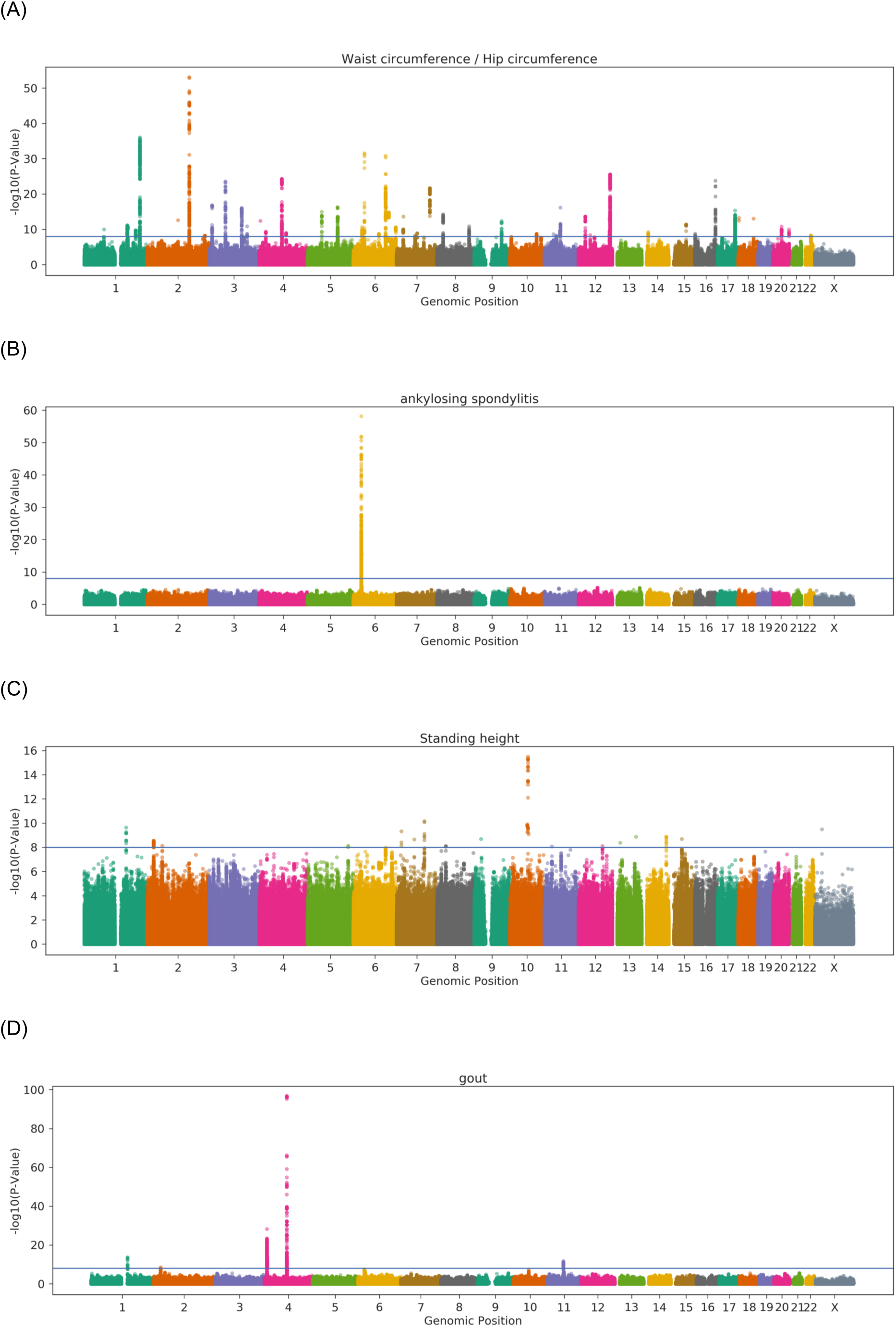
Manhattan plots for the traits with the most lead sdSNPs. The x axis corresponds to the genomic position in the genome, and the y axis the –log_10_ *p*-value of the statistical test for which the null hypothesis is that no difference between the sexes exists. Each point corresponds to a genetic variant. Points that go above the statistical significance line at –log_10_ *p* = 1×10^−8^ are considered to be sdSNPs. Traits represented: (A) waist-hip circumference ratio, (B) ankylosing spondylitis, (C) standing height, and (D) gout.

### Genome-wide genetic effect comparison between sexes across traits

We directly assessed whether each genetic variant in the genome had different effects in males and females through sex-stratified GWASs. Our genome and trait-wide genetic effect comparison between the sexes (see Methods) yielded a total of 61 (72.62%) non-binary and 42 (9.42%) binary traits with at least one autosomal genetic variant presenting a significantly different effect at a *p* < 1 x 10^−8^ threshold (Table S3), hereon termed as a sex-dimorphic SNP or sdSNP (Table 1 shows traits with the largest number of independent autosomal sdSNPs found). When testing the X-chromosome variants, we found 28 (33.33%) non-binary traits with at least one sdSNP. Considering the autosomal genome, the trait with the largest number of sdSNPs was waist-hip circumference ratio, a complex trait that has appeared frequently in analyses of sexual dimorphism in genetic architecture^12,19,32^. A total of 2,421 sdSNPs were found for this trait, which represent 100 unique loci after linkage disequilibrium (LD) clumping (see Supplementary Note). The trait with the most sdSNPs in the X-chromosome was hematocrit percentage, with a total of 12 that mapped to 5 unique loci after LD clumping (Table S1 and Table S3). Our results include replications of several previously reported loci for traits including anthropometric measurements or diseases like gout^19,21–23,32,33^.

**Table 1.**
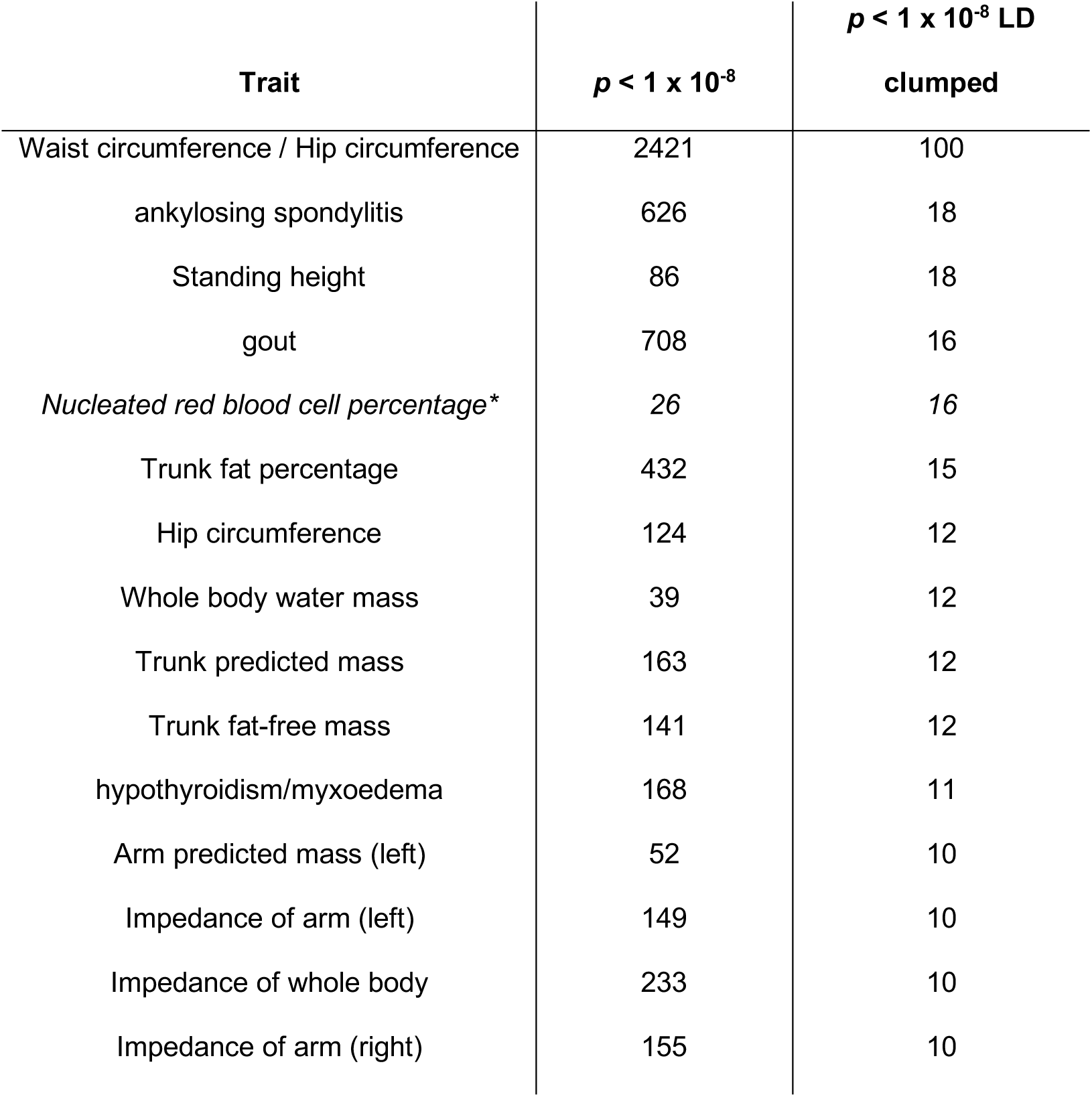
Traits with the largest number of autosomal lead sdSNPs. Sorted by number of loci post-LD clumping. (*) Our analyses point to nucleated red blood cell percentage likely being a false positive (see Supplementary Note).

A total of 4,179 (4,179/9,072,751 = 0.046%) and 4,196 (4,196/4,229,346 = 0.099%) autosomal genetic variants showed evidence of sexual dimorphism in at least one non-binary or binary trait respectively (*p* < 1 x 10^−8^), which mapped to 264 and 88 independent loci respectively. A total of 37 (37/7,227 = 0.213%) X-chromosome variants showed evidence of sexual dimorphism in at least one non-binary trait (*p* < 1 x 10^−8^), which mapped to 8 unique loci. The sdSNP associated to the highest number of traits (a total of 17) was rs115775278, an imputed intergenic variant found on chromosome 16. The closest genes to this variant include LOC105371341 (an uncharacterized non-protein coding RNA gene, with transcription start site, TSS, ∼40kb downstream), LOC390739 (MYC-binding protein pseudogene, TSS ∼50kb upstream), PMFBP1 (Polyamine Modulated Factor 1 Binding Protein 1, TSS ∼60kb downstream), and LINC01572 (a long intergenic non-protein coding RNA gene, TSS ∼470kb upstream). PMFBP1 has been linked to spermatogenesis function^34^. The distribution of hits across the autosomal and X chromosome genome is shown in Figure S3.

Several sanity checks were performed to support these results (see Supplementary Note, Methods), which included fitting alternative models, comparison to the Genetic Investigation of Anthropometric Traits (GIANT) cohort, and a randomization scheme. These checks suggested results for the nucleated red blood cell percentage trait likely represent false positives (see Supplementary Note).

### Phenotype prediction with sex-stratified estimates of effects

We studied whether genetic prediction could potentially be improved using sex-stratified models. To this end, we estimated genetic effects in a training population of 300,000 UK Biobank white British individuals in two different ways: (1) including both sexes in the model (obtaining sex-agnostic effects) and (2) using each sex in a separate model (obtaining sex-specific effects, see Methods). We then used a testing population consisting of 43,884 white British individuals to compare the performance of these two models in three different ways using PGSs: (1) obtaining predictions from the sex-agnostic effects (*agnostic* PGS), (2) obtaining predictions using the female effects applied to females and the male effects applied to males (*same* PGS), and (3) obtaining predictions using the female effects to predict in males and vice versa (*opposite* PGS). Prediction accuracy was measured as the correlation (r) between or the area under the curve (AUC) for our prediction and the true phenotype value for non-binary and binary traits respectively. Only lead sdSNPs were used in our PGS calculation. Due to the general low number of sdSNPs across traits, we focused our comparison on phenotypes with at least 10 lead sdSNPs. These included 7 non-binary traits (waist-hip circumference ratio, standing height, trunk fat percentage, hip circumference, whole body water mass, trunk predicted mass and trunk fat-free mass) and 3 binary traits (ankylosing spondylitis, gout and hypothyroidism/myxoedema).

Although of the 7 non-binary traits tested only waist-hip ratio showed a significantly different prediction accuracy (*p* < 0.05) between *same* PGS and *agnostic* PGS in males (Table S8), all 7 traits consistently presented a larger prediction accuracy when comparing the sex-stratified model with the agnostic model, thus suggesting that the stratified model captures the effect sizes better than the agnostic model. On the other hand, we consistently observed smaller prediction accuracies when the stratified model was used to perform predictions on the opposite sex (*opposite* PGS). We did not observe any consistent prediction improvements for the 3 binary traits considered (Table S8).

A limitation of our approach is the overlap between our discovery data set (used to establish sdSNPs) and our training and testing data sets in our prediction analysis (see Methods). We repeated our analysis with independent data sets (see Methods) for waist-hip circumference ratio, and we found that the *same* PGS and *agnostic* PGS had similar predictive ability in females (r = 0.132 with *p* = 1.85 x 10^−98^ and r = 0.133 with *p* = 4.02 x 10^−99^ respectively), the *same* PGS surpassing the *agnostic* for males (r = 0.038 with *p* = 7.91 x 10^−8^ and r = 0.024 with *p* = 9.97 x 10^−4^ respectively), however the differences in predictive power were not significantly different in either case (*p* > 0.05).

A possible explanation for the modest increase in predictive power found when using our sex-stratified models, when taking observed differences in heritability into account, is the potential existence of large numbers of SNPs of small dimorphic effect across the genome. These small effects remain undetected in a GWAS, and as such, are not being included in our predictions. This reasoning parallels the missing heritability problem^35^, where the predicted heritability of traits can’t be explained by the detected GWAS associations, a hypothesis for which is the existence of large amounts of variants of small effect that are yet to be found. Consistent with this theory, we found that our sdSNPs generally accounted for a very low percentage of the sex-specific heritability for the considered traits (Table S9, Methods), which ranged from 0.85% to 3.31%. Waist-hip circumference ratio was the exception, for which our sdSNPs accounted for 54.72% and 10.63% of the female and male specific heritability, respectively, which could be due to the substantially larger number of sdSNPs identified. This could also be, however, due to sdSNPs having a generally small effect on the phenotypes considered.

### Potential masking of loci of interest in sex-agnostic studies

Currently, many GWASs fit non-sex-stratified models. However, a situation could arise in which (1) a locus possesses a differentially signed genetic effect in each sex or (2) a genetic variant shows a larger effect in one of the sexes and a small or no effect in the other. In any of these situations, the power of detecting the variant will be reduced in a non-stratified analysis, and the variant effect size misestimated in both sexes. This phenomenon we term as “masking” of a genetic effect.

To assess masking effects in the UK Biobank, we evaluated the total number of genetic variants that were found to be significantly associated with a trait in a sex-stratified GWAS (i.e. associated to a trait in males and/or females), but were not significantly associated in the sex-agnostic model. We performed this analysis on the 530 traits in our study, considering a genetic variant as potentially masked if it is significant in females and/or males but not for the mixed population at a *p* < 1 x 10^−8^ threshold.

We found that 127/446 (28.48%) binary and 82/84 (97.62%) non-binary traits had at least one genetic variant that showed potential masking across the autosomal genome (Table S10, Figure S7A). On average, the percentage of these variants that presented opposite signs in each sex was 13.78% (SD 29.56%) in binary traits and 3.2% (SD 8.12%) in non-binary traits (Figure S7B). This may indicate that for a small percentage of traits opposite signed genetic effects are leading to masking. However, this could also be the result of smaller sample sizes leading to false positives in one sex but not the other.

We also found a significant correlation between the number of masked variants and the number of sdSNPs in both binary (r = 0.346, *p* = 0.042) and non-binary traits (r = 0.639, *p* = 3.916 x 10^−8^), as shown in Figure S8. On average, the percentage of masked variants that presented sexual dimorphism in binary traits was 2.45% (SD 13.48%), and 0.74% (SD 1.68%) in non-binary traits (Figure S7C). These low percentages could indicate that masked variants may have different effects on the two sexes, just not surpassing our significance threshold to be considered sdSNPs. On the other hand, 42 of our 103 traits with at least one sdSNP had one of these sdSNPs potentially masked and on average, the percentage of dimorphic variants that were potentially masked in binary traits was 12.30% (SD 30.59%) and 18.44% (SD 21.99%) in non-binary traits (Figure S7D). This could suggest a large number of potentially interesting variants that present a difference in genetic effect between the sexes could be being missed in sex-agnostic studies.

### Gene level analyses

To gain insight into the biological meaning of these results, gene enrichment analyses were carried out for all 103 phenotypes with at least one sdSNP. To do this, the two-tailed sex-comparison *p*-values for the sdSNPs found were converted to two one-tailed *p-*values (*p*_*F*_ and *p*_*M*_) according to the sex which presented the largest genetic effect (see Methods). Using these two sets of *p-*values in combination with MAGMA we then estimated the degree of sexual dimorphism of each gene, thus obtaining dimorphic gene lists, which were dominant in females or males (i.e. which presented a significantly larger effect in one sex versus the other, see Methods). The GENE2FUNC tool in FUMA was then used to investigate any functional enrichments among these dimorphic gene lists (Table S11 and S12, Supplementary Note) for the 10 traits with the largest number of dimorphic genes. These were largely all of the anthropometric class and were: standing height, waist-hip circumference ratio, trunk predicted mass, trunk fat-free mass, trunk fat percentage, whole body fat-free mass, basal metabolic rate, impedance of arm (left), body fat percentage and hip circumference. As a background to compare our results to, this procedure was repeated using sex-agnostic GWAS results, obtaining gene sets enriched in genes associated to each of our 10 phenotypes (see Methods).

A total of 4,840 gene sets were found to be enriched in either male or female dominant genes across the 10 traits considered (*q* < 0.05, Table S13, Table S15). Genes dominant in one sex or the other were found to be enriched in sets in an exclusive manner (i.e. sets would not show a larger amount of both male and female dominant genes than what would be expected by random), with the average percentage of shared enriched sets across traits being 3.3%, SD = 3.34% (Table S13). A total of 383/4,840 gene sets were found to be significantly differentially enriched between male and female dominant genes in at least one of the traits considered (Fisher’s exact test *q* < 0.05), with an average of 12.88% (SD 12.96%) of gene sets being dimorphically enriched across traits (Table S13). Furthermore, 251/383 gene sets were found to also show a significant difference in enrichment when comparing with the results of our background of sex-agnostic associated genes (Fisher’s exact test *q* < 0.05, see Methods), the mean percentage of gene sets presenting this behaviour across traits being 67.30% (SD 18.87%, Table S13).

Heatmaps were produced considering the aforementioned 251 gene sets, with hierarchical clustering both by gene set and by trait (Figure S10). Most notably, we find clusters of sets pertaining to small non-coding RNA (sncRNA) biogenesis and RNA-mediated silencing (Figure S10), enriched in female dominant genes for body mass-related traits. It has previously been postulated that miRNA may play a role in the regulation of phenotypic sexual differences due to their ability to regulate large numbers of genes with a high degree of specificity, with intervention of the sex chromosomes and/or gonadal hormones^36^.

### Sex differences in gene expression regulation

Differences between the sexes in complex traits could be partially explained by sex-specific gene expression regulation, which could lead to differences downstream across biological pathways and traits, and thus to the detection of GxS interactions in GWASs. Although studies have been carried out searching for differential gene expression between the sexes (sex-DE) in a variety of tissues of interest, studies linking sex to differences in gene expression regulation (sex-specific or sex-biased eQTLs) are few, with often contradictory results^5,37–40^. These mixed results could be due to the contribution of GxS to gene expression being tissue specific, a lack of sufficient statistical power, or the fact that this contribution occurs only on a small number of genes^39^. Overall, a system-wide analysis (i.e. across a wide variety of tissues) is yet to be carried out in order to determine whether there is evidence of sex-biased eQTLs and whether some tissues are more prone to sex-specific regulation than others.

In order to bring light to potential intermediary mechanisms underlying differences in genetic architecture between the sexes we investigated whether our lead sdSNPs could also be acting as sex-biased eQTLs. To do this, we performed an eQTL analysis, looking for GxS interactions in gene expression, considering genes within a 1Mb window to our lead sdSNPs (Table S3, see Methods). This was done for a total of 39 tissues from the Genotype-Tissue Expression (GTEx) consortium v6, originating from up to 450 individuals.

A total of 8 sex-biased eQTLs were found at a *q* < 0.05 threshold (Table S16, S17 and S18). We also checked for enrichment of GxS in gene-variant pairs for variants that presented evidence of sexual dimorphism (genetic effect comparison between the sexes *p* < 1 x 10^−8^) versus those that did not (genetic effect comparison between the sexes *p >* 0.5), using contingency tables (see Methods). We found enrichment for a small number of the tissues considered (see Supplementary Note).

The variant rs56705452 from chromosome 6 was found to be a sex-biased eQTL for the transcript ENSG00000204520.8, which corresponds to the gene MICA (Figure S11), in muscle tissue. This gene encodes the highly polymorphic major histocompatibility complex class I chain-related protein A, and variations of this gene have been associated to susceptibility to psoriasis, psoriatic arthritis and ankylosing spondylitis, amongst others^41^. Interestingly, this sdSNP has been shown to bind FOXA1 through ChIP-Seq experiments^42^, a protein that dictates the binding location of androgen and oestrogen receptors, and that has been found to play a role in the sexually dimorphic presentation of various cancers^43,44^. Furthermore, this sdSNP was found to act in a sexually dimorphic manner in regards to its association to ankylosing spondylitis in our genome-wide sdSNP analysis. This result would be consistent with a hypothesis where this sdSNP is regulating MICA in a sex-dependent manner in the muscle tissue, thus leading to differences in ankylosing spondylitis presentation between the sexes when one possesses a particular variant.

The small number of significant sex-biased eQTLs found parallels the findings of recent study by Porcu et al^40^, where they conclude that millions of samples would be necessary to observe sex-specific trait associations that are fully driven by sex-biased eQTLs. Overall, this type of pipeline could help in the future when short-listing biomarkers for risk susceptibility in men and women, help develop precision medicine strategies for each of the sexes, and bring light into the underlying mechanisms of the disease/trait of interest as well as possible underlying sexually different molecular networks, once larger sample sizes become available.

## Discussion

In this study, we have delved into the differences in genetic architecture between the sexes in the UK Biobank for a total of 530 traits from around half a million individuals. This has enabled us to assess the genetics of sexual dimorphism at a depth and breadth not previously achieved.

Overall, we have found evidence of dimorphism for a large number of the traits considered, though be it of generally modest magnitude, through a thorough investigation of sex-specific genomic parameters. A total of 71 traits were found to present significantly different heritability estimates between the sexes, while a total of 69 presented genetic correlations between the sexes that significantly differed from one, indicating the presence of genetic heterogeneity across these complex traits. In order to dissect this heterogeneity and pin-point genetic sites that could be dimorphically associated to these traits, sex-stratified GWASs were performed, yielding over 100 traits with at least one sdSNP. These traits included those of the anthropometric class as well as diseases like gout, ankylosing spondylitis or hypothyroidism. These results reinforce the need for future studies to account for genetic sexual heterogeneity to fully understand the genetic underpinnings of disease and ultimately shed light on potential sex-specific biological mechanisms.

Having found evidence of GxS across the genome, we investigated whether sex-specific genetic models could improve phenotypic prediction. While a significant improvement in prediction was only found for one of the traits considered, a consistent trend of increased predictive accuracy was seen when comparing the results of sex-specific models to those of a sex-agnostic model. Putting our results in context with the heritability differences found, we postulate the potential existence of large numbers of loci presenting small amounts of dimorphism with insufficient statistical power to be detected in our analysis that could account for both this missing heritability difference as well as the absence of increased predictive power. We also investigated whether sex-agnostic models could potentially be missing loci of interest, and found indications of potential masking for 209 traits, with further investigation being needed to replicate these results.

Finally, gene set enrichment and eQTL analyses were performed in an effort to translate our GWAS results to function and bring light to potential mechanisms underlying the observed dimorphism across traits. Our eQTL analysis found a total of 8 sex-biased eQTLs, but our results parallel previous reports on the need for larger sample sizes to truly uncover potential links between sex-biased eQTLs and sdSNPs. Our gene-set enrichment analysis suggests a link to miRNA regulation, which has been hypothesized in the past to underlie sexual dimorphism. Further studies are needed to truly understand what underlies sdSNPs, moving beyond gene expression regulation mechanisms and looking at other biological regulatory mechanisms and omics data sets.

## METHODS

### UK Biobank data

UK Biobank is a large population-based prospective study with participants aged 40 to 69 years at recruitment, with extensive matching phenotypic and genomic data^45^. In this study, of the circa 490,000 individuals whose data was released in July 2017, we considered data pertaining to a total of 452,264 white European individuals (245,494 females and 206,770 males) after excluding individuals that were identified by UK Biobank as outliers based on genotyping missingness rate or heterogeneity and individuals whose self-reported sex did not match that inferred from their genotypes. We also removed individuals whose first or second genomic principal component differed by over 5 standard deviations from the mean of self-reported white Europeans. Finally, we removed individuals with a missingness rate > 5% for the genetic variants that passed quality control (described in Genotypes section), arriving at the aforementioned number of individuals.

#### Genotypes

UK Biobank’s participants were genotyped using either of two arrays, the Affymetrix UK BiLEVE Axiom or the Affymetrix UK Biobank Axiom array, and later augmented by imputation of over 90 million genetic variants from the Haplotype Reference Consortium, the 1000 Genomes project, and the UK 10K project.

We excluded variants which did not pass UK Biobank quality control procedures in any of the genotyping batches and retained only bi-allelic variants with *p* < 1×10^−50^ for departure from Hardy-Weinberg and MAF > 1×10^−4^, computed on a subset of 344,057 unrelated (Kinship coefficient < 0.0442) individuals of White-British descent with missingness rate > 2% in the study cohort, paralleling the QC procedure followed by Canela-Xandri et al^46^.

A second round of QC was done prior to performing everything that followed the sex-stratified GWAS and genetic parameter estimation (described below), including calculation of genetic correlations and beta comparisons between the sexes. This was done to account for the smaller sample sizes (versus those of ^46^) as well as to be more stringent as we are comparing between groups. Variants were further filtered if they possessed *p* < 1 x 10^−6^ for departure from Hardy-Weinberg and MAF > 1 x 10^−3^, computed on the aforementioned subset of unrelated individuals. A stricter MAF threshold (MAF > 1 x 10^−1^) was set for binary traits due to the general limited number of cases available which can lead to an inflation of type I error rates in association tests^47^. Furthermore, variants with a significant effect (*p* < 1 x 10^−8^) when running a GWAS on sex for the aforementioned subset were also excluded, as these could arise due to sampling bias^48^. Finally, only imputed variants with no genotyped counterpart and with an imputation score > 0.9 were retained. As a result, a total of 9,072,751 (602,984 genotyped and 8,469,767 imputed) autosomal genetic variants and 17,364 X-chromosome genetic variants remained for our analysis of non-binary traits, and 4,229,346 autosomal (244,743 genotyped and 3,984,603 imputed) and 7,227 genotyped X-chromosome genetic variants for our analysis of binary traits.

#### Phenotypes

In total we analyzed 530 non-sex specific traits. These included 446 binary traits, which had at least 400 cases in each of the sexes, relating to self-reported disease status, ICD10 codes from hospitalization events, and ICD10 codes from cancer registries, as well as 84 non-binary traits comprising non-scale transformed continuous and ordered integral measures. For more information on the treatment and filtering of the phenotype data see ^46^.

### Sex-stratified parameter estimation

To obtain genetic and environmental variance estimates for each of the sexes we used DISSECT^28^ following the same methodology as described in ^46^. Briefly, a partition of variance analysis was run using linear mixed models (LMMs), which were fitted for each trait with a Genetic Relationship Matrix (GRM) containing all common autosomal genetic variants (MAF > 5%) which passed QC.

Heritability was then calculated as

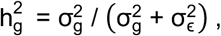

where 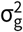 and 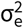 are the estimates of the genetic and residual variance. Heritability for binary traits was transformed from the observed scale to the liability scale using the sex-specific population prevalence of the trait, under the assumption that there was an underlying normal distribution of liability to the considered trait, as described in ^49^.

Wanting to see if the sexes differ in regards to their ability to undergo adaptation, evolvability was calculated for males and females separately. Evolvability, defined as the expected evolutionary response to selection per unit of selection^50^, or the ability of populations to respond to natural or sexual selection^51^, was calculated as

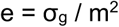

where σ_g_ is the additive genetic variance of the trait and m is the trait mean.

To establish differences between heritability, genetic variance, and evolvability between the sexes, we used the t statistic

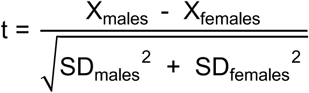

where X represents either heritability, genetic variance or evolvability, assumed to be independent between the sexes, and SD the standard deviation of the aforementioned, for males and females respectively. *p*-values were then FDR corrected (using the Benjamini-Hochberg procedure) to account for multiple testing.

### Sex-stratified GWAS

To test the genetic effect of each variant for each of the sexes on the 530 chosen traits we ran a sex-stratified GWAS. The procedure followed for each of the sexes is that described in ^46^ using the DISSECT software^28^. Briefly, a Linear Mixed Model (LMM) was run,

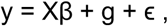

y being the vector of phenotypes, X the matrix of fixed effects, β the effect size of these effects, g the polygenic effect that captures the population genetic structure, and ϵ the residual effect not accounted for by the fixed and random effects. Following the procedure described in ^46^, our curated genetic variants were regressed against the residuals of the LMM to assess association.

### Genetic correlations

Genetic correlations between the sexes were calculated using the bivariate linkage disequilibrium score (LDSC) regression analysis software^30^, which works directly on GWAS summary data and can thus be applied to very large sample sizes. As we were using data of European origin, we used the LD scores provided by the LDSC software and limited our genetic correlation calculation to the genetic effect estimates of SNPs for which such scores were available (1,189,831 total genetic variants, of which 1,169,868 passed LDSC’s QC filters and were used in the computation). These scores were computed using the 1000 Genomes European data. We furthermore restricted our binary traits to those that had at least 5000 cases in each of the sexes, as was recommended in the documentation. Note that for traits with very low heritability this computation was unsuccessful. In total, genetic correlations were obtained for 83 non-binary and 77 binary traits.

To establish which correlations differed from one, we used the t statistic

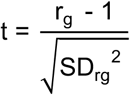

where r_g_ is the genetic correlation, and SD_rg_ is the standard deviation of the genetic correlation. FDR correction was applied to account for multiple testing.

### Sex differences in genetic effects

To compare genetic effects across the genome between the sexes for all traits we considered the following t-statistic, as used in previous sex-stratified GWAS comparison studies^19,22^:

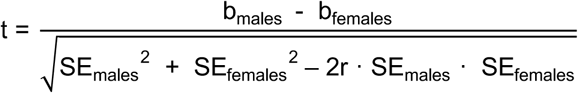

where b is the estimated effect of the genetic variant considered for a given trait for males and females, SE is the standard error of the effect, and r is the Spearman rank correlation between the sexes across all genetic variants for a given trait. Both the SE and b were adjusted by the standard deviation of the trait, for each sex, to correct for scale effects that could act as confounders in the study. Some studies have opted to ignore the third term in the denominator, which estimates the covariance of the error terms, assuming r to be equal to 0^6^. However, for the traits considered this correlation ranged from - 0.00335 (cervical spondylosis) to 0.34173 (standing height), thus our decision to include it.

As this test was done for all variants and all phenotypes, this effort resulted in a total of 4,808,558,030 statistical tests (9,072,751 x 530). Binary traits were then filtered further as stated in the Genotypes section. To account for multiple testing, we considered the commonly used genome wide significance cut-off of *p* < 1×10^−8^.

In order to cluster our results into independent lead variants, we used the --clump option in PLINK 1.9^52^. For each individual trait, variants found to be genome-wide significant with regards to difference between the sexes were clustered into lead variants, assigning them variants in LD within 10Mb, with an r^2^ > 0.2 with the lead variant. To obtain the total number of independent loci across all traits, the same clustering method was used but for all variants found to be leads across traits, choosing the variant with the lowest *p*-value if variants were found in more than one trait.

### Analysis checks

Our replication/sanity check methods can be divided into technical (different models, randomization) and biological (comparison to the GIANT, cohort). Detailed results of these analyses can be found in the Supplementary Note.

#### Different models to check for dimorphism

Using a subset of our original data of the UK Biobank’s 344,057 unrelated White British participants (Kinship coefficient < 0.0442), of which 158,956 are males and 184,928 are females, we proceeded to run two new models with which to contrast our original results for our candidate sexually dimorphic genetic variants. The models looked to replicate our original model in all but a few aspects, keeping all covariates the same. The first (Model 1) included a gene by sex interaction term (GxS), thus allowing for the running of males and females simultaneously. The second model (Model 2) consisted on the inverse rank normalization of non-binary phenotypes within sex prior to performing an identical sex-stratified GWAS to the original. Furthermore, Model 1’s phenotypes were normalized by the standard deviation within trait and sex, and Model 2’s binary traits also followed the same treatment. We considered methodological replication occurred for Model 1 when the GxS interaction term was significantly different from 0 considering different significance cut-offs, as stated in the Results section. For Model 2 we performed the same statistical test as we originally had when testing for differences in genetic effects, considering replication at difference significance cut-offs as well. These models were run using the PLINK software^52^.

#### GIANT comparison

GIANT sex-stratified anthropometric European summary statistics from 2015^53^ were downloaded for waist circumference, hip circumference, and waist-hip ratio from the GIANT portal (https://portals.broadinstitute.org/collaboration/giant/index.php/GIANT_consortium_data_files).

Summary statistics for height and weight were also downloaded from their 2013 dataset, also derived from a European population^22^. These were used to perform the same statistical test done originally to check for differences in genetic effects.

We also compared our Waist-hip ratio results to the more recently published GIANT-UK Biobank meta-analysis for human body fat distribution^20^, for which the summary statistics were downloaded from https://github.com/lindgrengroup/fatdistnGWAS. Again, taking their summary statistics we performed the same statistical test as we did with our data to check for differences in genetic effects.

#### GWAS with randomized sex

As a further means of obtaining evidence to support our results we repeated our methodology on randomized sex samples, expecting, given the assumption that no unknown factor correlated with sex was still present, that dimorphism would not appear beyond what would be expected by chance. This randomization consisted of assigning males and females to two groups randomly, and then checking for genetic effect differences between the two. The proportion of males and females in UK Biobank was preserved within the groups. We limited this analysis to chromosomes 1 and 6, across all of our 530 traits, due to these chromosomes showing large numbers of GWAS hits in ^46^.

In our original analysis, our association tests were performed in two stages following the methodology described in ^46^, first running phenotypes against all covariates and the polygenic effect (Step 1), and then regressing genetic variants against the residuals to assess association (Step 2). Whereas in our original model Step 1 was run separately for males and females, thus obtaining sex-specific residuals, here Step 1 was run with both males and females, thus obtaining a sex-agnostic residual estimate. This was then used to obtain genetic effects across chromosomes 1 and 6 for our randomly assigned groups, hereon termed Group 1 and Group 2.

As in our original analysis we used sex-specific residuals, the results of Group 1 and Group 2 are not directly comparable. To overcome we also re-ran the analysis for males and females using the sex-agnostic residuals for chromosomes 1 and 6, which we then used to compare with our randomization results.

To check for differences in genetic effects between Groups 1 and 2, the same methodology described previously was used, where instead of standardizing genetic effect and standard error estimates by the standard deviation of the phenotype within sex, this was analogously performed using the standard deviation of the phenotype within the random groups, in order to keep the methodology as similar as possible.

### Calculation of Polygenic Scores (PGS)

Using again the subset of unrelated White British individuals from UK Biobank, we calculated PGS for our traits with most evidence of dimorphism found using our lead sdSNPs.

PGS are calculated as:

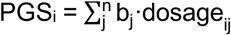

Where PGS_i_ is the polygenic risk score estimate for individual i, b_j_ is the genetic effect estimate for variant j, dosage_ij_ is the effect allele count (0, 1 or 2) of variant j in individual i, and n is the total number of genetic variants considered in the PGS calculation (here just the lead sdSNPs).

The genetic effects used in our PGS calculations were obtained re-running our original model using UK Biobank’s unrelated White British participants, randomly selecting 150,000 women to obtain female specific genetic effects, 150,000 men to obtain male specific genetic effects, and 75,000 men and 75,000 women to obtain sex-agnostic genetic effects. This was done to match sample sizes.

We then proceeded to calculate PGS for a total of 34,928 women and 8,956 men from UK Biobank and which were self-reported white European, which had not been considered in the calculation of the genetic effects.

We did this in three ways: using the genetic effects corresponding to the sex of the individual (*same* PGS), using the genetic effects corresponding to the opposite sex of the individual (*opposite* PGS) and using the genetic effects for the whole population (*agnostic* PGS). Thus, in total each of our circa 44K individuals had 3 PGS calculated.

In order to assess predictive power, the phenotypes of our 44K individuals (corrected by all the covariates in our original model to account for population structure and other effects) were then either regressed on the *same, opposite* and *agnostic* PGSs respectively in the case of non-binary traits, and in the case of binary traits the area under the curve was calculated for the ROC curve for the three PGS groups.

An important caveat of this methodology is that there is an overlap between the discovery (the population used to declare variants as sexually dimorphic) and the replication (for which genetic effects were re-calculated and/or PGRS were obtained) populations. As a way of checking whether the overlap could potentially be influencing our results, we repeated this step by obtaining the genetic effects for the circa 408K individuals of White British ethnicity in UK Biobank, repeating the steps described in the “Sex differences in genetic effects” section, to establish genetic effect differences across the genome. Then we calculated the different PGSs for the remaining circa 42K individuals of white ethnicity, again regressing on the phenotype, for our most representative trait: waist-hip circumference ratio. This, however, only serves as a validation for a single trait, meaning that caution should be taken when interpreting the predictive power of the PGSs calculated.

### Heritability explained by sdSNPs

In order to put our prediction analysis into context, we obtained the proportion of sex-specific heritability explained by the sex-specific genetic effect estimates of the sdSNPs found for each trait. To do this, for each trait with i sdSNPs, the heritability of sdSNPs was calculated as

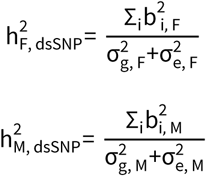

where b_F_ and b_M_ are the sex-specific genetic effects for each of the i dsSNPs for a given trait, and where 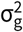 and 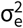 are the sex-specific genetic and residual variance estimates for each trait respectively. The proportion of the sex-specific heritability explained by sdSNPs was then calculated for each sex as 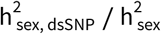.

### Gene level analysis

We performed a gene-level analysis to translate our SNP data into a more manageable and interpretable form. This was done for the subset of traits that presented at least one sexually dimorphic variant for a total of 103 traits.

As we wanted to obtain genes relevant to each of the sexes, we began by partitioning our two-tailed *p*-values (*p*_2T_) from the genetic effect (b) comparison between the sexes into two one-tailed *p*-values. For genetic variants where b_F_ was greater than b_M_, one tailed *p*-values were calculated as:

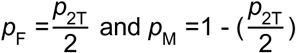

On the other hand, when the beta was greater in males, the *p*-values were calculated as

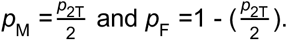

This process led to the creation of two additional distinct sets of *p*-values for each phenotype, corresponding to sites where the genetic effect was significantly greater in males or females.

Each of these sets of *p*-values (*p*_2T_, *p*_M_ and *p*_F_) were subsequently used to identify gene level associations using MAGMA^54^. First, we annotated every gene (i.e. defined which SNPs were in the gene region) considering a range of 1kb upstream and downstream. MAGMA was then run for each phenotype and each set of one tailed *p*-values separately, considering two distinct SNP-wise models (SNP mean and top SNP), using a random sample of 1,000 unrelated white British individuals from the UK Biobank, 500 males and 500 females, as a base population for LD and MAF correction. The analysis provides three distinct *p*-values for each gene, one for the SNP mean model, one for the SNP top model, and a combined *p*-value. For subsequent analyses, we considered the combined *p*-value for each gene. Genes were declared significantly dimorphic and female or male dominant if an FDR corrected combined *p*-value < 0.01 was obtained when considering *p*_*F*_ or *p*_*M*_ respectively.

The set of genes that reached our threshold were then used in the gene set and enrichment analyses using the GENE2FUNC tool in FUMA^55^. The GENE2FUNC tool in FUMA was run for the top 10 traits with the largest number of significant genes when considering a two-tailed *p*-value. As a result, FUMA was run for 10 traits for both male and female dominant genes.

Briefly, FUMA takes our list of candidate genes and checks for enrichment across: (1) differentially expressed genes (DEG) for different tissues (tissue enrichment analysis was conducted considering the GTEx V6 database^56^), and (2) biological pathways/functional categories (considering MSigDB v7, WikiPathways and GWAS Catalog^57^). Enrichment is assessed using a hypergeometric test. In this study we have focused on just biological pathways/functional categories. Significantly differentially enriched gene sets between female and male dominant genes were assessed using a Fisher’s exact test, and *p*-values were FDR corrected.

As a background, the same procedure as is described in this section was followed using sex-agnostic GWAS results for the 10 traits of interest. Using MAGMA, genes associated to each of the traits were obtained, and sets enriched for genes associated to each trait were obtained using FUMA. Significantly differentially enriched gene sets between our background and our female and male dominant genes were obtained using a Fisher’s exact test, and *p-*values were FDR corrected.

Heatmaps of the scaled -log10 *q-*values per set were created for each phenotype and sex. Heatmaps for gene sets were limited to those both significantly differentially enriched in at least one sex versus our background (Fisher’s exact test *q* < 0.05), as well as significantly differently enriched between female and male dominant genes (Fisher exact test *q*-value < 0.05).

### GTEx data

In order to assess whether differences in genetic architecture, as established by our analysis of the UK Biobank data, lead to differences in gene expression, we proceeded to complete an expression quantitative trait loci (eQTL) analysis looking for gene x sex interactions in gene expression. The data from the Genotype-Tissue Expression Project (GTEx) V6p release was used, which consists of samples and genotypes from 449 human donors (292 males and 158 females) and 39 non-sex specific and non-diseased tissue types. Each tissue type holds a different number of samples (minimum of 70, median of 149), with a male-bias present in all of them (the percentage of female samples ranging from 25% to 44%).

Processed, filtered and normalized RNA-seq data was downloaded from the GTEx portal for both the autosomal genome as well as for the X chromosome, the number of transcripts varying across tissues due to tissue-specific expression, with a median of 23,538 autosomal transcripts and 793 X-linked transcripts per tissue. Covariates were also downloaded for each of the considered tissues. Further information regarding the treatment of the data can be found in the GTEx v6p analysis methods documentation.

Genotype data was also obtained for all the donors from dbGaP, and which included a total of 11,607,846 genetic variants, from genotyping efforts with Illumina OMNI 5M and 2.5M SNP Arrays and imputation from the 1000 Genomes Project Phase I version 3 reference panel. For more information regarding the processing of the genotype data see the GTEx v6p analysis methods documentation.

### eQTL analysis

To assess whether our dimorphic genetic variants had different effects on the expression of nearby genes depending on sex, we assigned our lead sdSNPs nearby genes (1Mb window) using *Granges*^58^ and the Biomart resource^59^. The total number of sdSNP-gene pairs were 6591 when considering our autosomal hits for non-binary traits, 4533 when considering our autosomal hits for binary traits, and 95 when considering our X-chromosome hits for non-binary traits.

For each of the 39 tissues, we tested for GxS in gene expression for each variant-gene pair using a linear regression model in PLINK 1.9, and which was adjusted for three genotyping principal components (PCs) and PEER factors, the number of which included in the model depended on the sample size (sample sizes < 150 had 15 PEER factors, sample sizes between 150 and 250 had 30 PEER factors, and sample sizes over 250 had 35 PEER factors), as indicated in ^60^. In the end, our gene expression model for each gene within 1Mb of a sexually dimorphic variant, and each tissue was formulated as

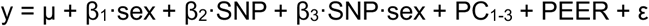

where y is the gene expression of the given gene in a given tissue, μ is the mean expression levels, β_1_ and β_2_ are the regression coefficients for sex and genotype of the sexually dimorphic variant respectively, β_3_ is the regression coefficient for the interaction of the genotype with sex, PC_1-3_ and PEER are the principal components and PEER factor covariates, and ε is the residual. FDR correction was applied to account for multiple testing.

When running our eQTL analysis we found that two tissues returned missing values across all tests performed with PLINK, the brain anterior cingulate cortex BA24 and the small intestine terminal ileum tissues. These are the two tissues with the smallest number of samples, therefore this absence of results is likely due to not enough variation being present in the phenotype.

### eQTL enrichment

To assess whether our sexually dimorphic genetic variants were enriched for GxS interactions versus those not presenting sexual dimorphism, we proceeded to run an exact replica of our eQTL model using genetic variants significant for the whole population (p < 10^−8^) but that had no evidence of being sexually dimorphic (t-statistic comparing genetic effects between the sexes with *p >* 0.5). Using contingency tables and Fisher’s exact test, we considered whether the number of significant variant-gene GxS terms was enriched for our sexually dimorphic variants for each of the tissues.

## ACKNOWLEDGEMENTS

This research was funded by the BBSRC through programme grants BBS/E/D/10002070 and BBS/E/D/30002275, MRC research grant MR/P015514/1, and HDR-UK award HDR-9004. This research has been conducted using the UK Biobank Resource under project 788.

## AUTHOR CONTRIBUTION

A. Tenesa conceived the study. A. Tenesa, O. Canela-Xandri, K. Rawlik and E. Bernabeu designed the genetic architecture, prediction, masking and eQTL analyses. A. Talenti and J. Prendergast designed the gene-level analyses. O. Canela-Xandri, K. Rawlik and E. Bernabeu pre-processed the data and conducted modelling. E. Bernabeu conducted the statistical analyses and prepared the initial manuscript. All authors contributed and commented on the development of the manuscript.

## CODE AVAILABILITY

We used DISSECT (v1.15.2c, May 24, 2018), which is publicly available at http://www.dissect.ed.ac.uk/ under GNU Lesser General Public License v3. We also used PLINK (v1.9 and v2.0, freely available online), BGENIX (v1.0 freely available online at https://bitbucket.org/gavinband/bgen), LD Score Regression (v.1.0.1, freely available online at https://github.com/bulik/ldsc), MAGMA (v1.06, freely available online at https://ctg.cncr.nl/software/magma) and FUMA (freely available online at https://fuma.ctglab.nl/). Custom code created to perform the analysis is openly available from the University of Edinburgh DataShare repository [DOI’s to follow].

## DATA AVAILABILITY

This research has been conducted using the UK Biobank Resource under project 788. The protected data for the GTEx project (for example, genotype and RNA-sequence data) are available via access request to dbGaP. Processed GTEx data (for example, gene expression and eQTLs) are available on the GTEx portal: https://gtexportal.org. The authors declare that the data supporting the findings of this study are available within the paper and its supplementary information files. The GWAS summary statistics are openly available from the University of Edinburgh DataShare repository [DOI’s to follow].

