## Supplementary Note for "Sexual differences in genetic architecture in UK Biobank"

#### Waist-hip circumference ratio

Waist-hip circumference ratio is a complex trait that has frequently been of interest in sexual dimorphism studies, and for which the largest number of independent sdSNPs have been reported, Winkler and collaborators citing 44 in 2015[1], and a recently published meta-analysis including both the GIANT consortium and UK Biobank's data reporting 105[2]. Here we report 100 independent sdSNPs at a $p < 1x10^{-8}$ threshold, a smaller number likely due to the smaller sample size and stricter significance threshold (a more detailed comparison of these sdSNPs is found in the "Comparison with GIANT" section of this Supplementary Note). Our results corroborate previously published sdSNPs nearby genes including COBLL1, VEGFA, LYPLAL1, RSPO3, CMIP, ADAMTS9, FAM13A, and CMIP amongst others. Figure 3A shows the distribution of sdSNPs for waist-hip circumference ratio across the genome.

When looking closer at our 100 lead sdSNPs, we find that 66 show opposite effect directions in males versus females. Considering the remaining 34 sharing effect direction, 20/34 have a greater effect in females. This dimorphism can be seen in Figure S4A, where female genetic effects are plotted against male genetic effects, and where dimorphic SNPs are colored in red. It is easy to see a clear divergence from the x = y slope, with most genetic effects in males being close to 0 as opposed to the females', who possess larger absolute effects. Furthermore, of these 100 lead sdSNPs we find that 80 are significant in females at a $p < 1 \times 10^{-8}$ threshold, while only 6 loci are significant in males. A *p*-value comparison for all genetic variants across the genome is shown in Figure S4B, where sdSNPs are shown in red (note that all variants across the genome are shown, not just independent loci). In this plot, a large number of variants showing small *p*-values in females are shown to have larger counterparts in males.

#### Analysis checks

To garner further evidence into dimorphism in genetic architecture we looked to support our results both technically (using different models and through a randomization scheme) and biologically (comparing our results to the GIANT consortium's).

##### *Comparison with GIANT*

We found that our results correlated with those from GIANT[1] for the available traits (waist-hip circumference ratio, hip circumference, waist circumference, standing height and weight), noting the smaller sample size, and that total overlap of our declared dimorphic hits was not found (i.e. we declared 100 leading SNPs/hits as dimorphic for waist-hip circumference ratio, and data was only available for 39 of these in GIANT, Table S4). We took GIANT's sex-stratified genetic effect estimates and tested them for sexual dimorphism, exactly as we had done with our own and as is described in the Methods section. Waist-hip circumference ratio had the largest correlation between GIANT's genetic effect comparison *p*-value and our own, likely due to its large dimorphism being detected with smaller sample sizes (Pearson and Spearman correlation being 0.84 and 0.62 respectively), followed by standing height (Table S4). The remaining traits available in GIANT showed similar behaviors. Significant correlations were also found when comparing GIANT's to our own female and male genetic effects.

As of early 2019, Pulit and collaborators have published a meta-analysis for the distribution of human body fat[2]. This study includes both the GIANT consortium data as well as the UK Biobank data, totaling up to the biggest sample size seen in a study of this nature, with 694,649 individuals of White European ascent. Their study also includes a sex-stratified analysis of Waist-hip circumference ratio, the results of which we have compared to our own. Pulit et al declared 105 independent loci as showing sexual dimorphism for waist hip ratio. Albeit our studies use differing methodologies (such as the pre-processing of phenotypes, or our significance cut-offs for declaring dimorphism), we find extremely good replication for both our declared sdSNPs as well as theirs. Figure S5A compares their genetic effect estimates for each of the sexes for our declared 100 lead sdSNPs, while Figure S5B compares the same but for their declared 105 lead sdSNPs.

To ease the comparison between our study and Pulit et al's, we used their summary statistics to obtain sdSNPs the same way we had done with our own data, thus establishing a new set of sdSNPs from their study (a total of 2,846 hits at $p < 1 \times 10^{-8}$ significance cut-off, without LD clumping), for which genetic effects were also compared with our own in Figure S5C. Finally, we compared the *p*-values for the dimorphism test of the latter to our own, again finding a high correlation despite many of these SNPs not being established as dimorphic in our own study due to our significance cut-off (Figure S5D). The fact that at the same threshold but with increased sample size more dimorphic hits (2,846 vs 2,418, pre-LD clumping) are found might suggest that once more samples become available, more dimorphism will be found for both this trait as well as others.

##### *Different models*

Our technical validation steps included the running of a linear model with a GxS interaction term (Model 1), testing for association of each phenotype-sexually dimorphic variant pair, as well as a sex-stratified linear model where the phenotypes were inverse rank normalized within sex prior to association testing (Model 2), similarly to Winkler et al[1] and other GIANT publications. We found good replication for both models, all traits showing at least 80% replication considering a $q < 0.05$ threshold, except nucleated red blood cell percentage in Model 2 (Table S5). This trait presents a very low heritability estimate in both sexes, only being significantly heritable in males (Table S1). Furthermore, when calculating the proportion of heritability that our sdSNPs account for, we find that this estimate surpasses 1, indicating that these SNPs are likely false positives and that their genetic effects are not accurate. Because of this, this trait has been discarded from future discussion. For the remaining traits, given the same threshold as was originally used for assessing sexual dimorphism ($p < 1x10^{-8}$) is used in our replication stage, it is not surprising that the number of sdSNPs found is lower than in our original analysis, as a smaller subset of the original UKB cohort was considered.

##### *Randomization*

As a further means of validation, we repeated our analysis through a randomization scheme (see Methods). To do this, we took our circa 450K individuals and assigned them randomly to two groups, Group 1 and Group 2, thus creating an alternative to our original two groups defined by sex, males and females. We then proceeded to re-calculate the genetic effects across chromosomes 1 and 6 (for which we found a large number of sdSNPs in our original analysis) for the new randomized groups and repeat our test checking for significant difference in genetic effects at a $p < 1x10^{-8}$ threshold.

As described in the Methods, we repeated our GWAS on the randomized groups using the residuals that had been estimated for a sex-agnostic fitting of the phenotypes against the covariates, as opposite to the sex-specific fitting that we had performed in our original study. Thus, to make sure that this wasn't altering our results, besides calculating genetic effects for our new random groups, we also proceeded to re-calculate the genetic effects for males and females using the sex-agnostic residuals. In this section we will refer to our original results as "Original", while the new sex comparison will be termed "Females vs Males" and the results for our randomized groups will be termed "Group 1 vs Group 2".

The Females vs Males comparison replicated our Original analysis very well (with an average percentage of 80.9% with SD 35.6% of our original dimorphic variants also being found in the new analysis at a $p < 1x10^{-8}$ threshold), while the Group 1 vs Group 2 on average showed no sexual dimorphism for both chromosomes and across traits (Table S6). QQ plots were also obtained to compare the distribution of *p*-values for the Females vs Males and the Group 1 vs Group 2 analyses, and a clear deviation from the null hypothesis is seen across all traits considered for the former but not the latter (shown in Figure S6A is malabsorption/coeliac disease, the trait with the largest amount of dimorphic hits pre-LD clumping in chromosomes 1 and 6).

We observed a larger number of sexually-dimorphic hits being found at the same significance threshold in the Females vs Males analysis than in our Original analysis for several traits, so to make sure that these signals corresponded to the same loci and were just the result of inflated *p*-values of nearby SNPs in LD we obtained Manhattan plots comparing the Female vs Male analysis to our Original analysis, determining that indeed this was the case. The Manhattan plot comparing these results for hypothyroidism, which presented 146 dimorphic SNPs in our original analysis and 793 in the Females vs Males analysis, is shown in Figure S6B. A potential reason behind the inflated *p*-values found in the Females vs Males analysis versus our Original analysis could be the effect of reporting bias, as described by Pirastu et al[3], where the authors state that including sex as a covariate in GWAS studies could bias effect estimates of individual variants if reporting bias is present in a given cohort.

#### Behaviour of hits within sex

We decided to investigate further how each of our found dimorphic loci behaved within the sexes in order to elucidate possible common characteristics or trends within them (i.e. did these loci generally present opposite sign genetic effects in males versus females, are these loci significant in just one of the sexes etc).

We found that the average percentage of hits in non-binary traits that are significant in a GWAS considering just females ($p < 1 \times 10^{-8}$) is 24.12% (SD 27.39%) and considering just males 29.69% (SD 30.84%), these numbers changing to 55.46% (SD 47.32%) in females and 66.09% (SD 44.17%) in males for binary traits. We also see that the majority of these sdSNPs present opposite genetic effect signs across non-binary traits (average 75.44%, SD 29.11%), the number decreasing to 35.57% (SD 45.10%) for binary traits. Finally, of the sdSNPs that were found to have the same effect sign between the sexes, on average 68.76% (SD 37.06%) of them presented a greater effect in females than in males in non-binary traits, while for binary traits the average was 47.76% (39.67%).

#### Sex-stratified versus cohort-wide analysis

We next proceeded to see how frequent sexual dimorphism is across the sdSNPs that are found in a mixed-sex population cohort, or, in other words, how many of the genetic variants that are found to be associated to a trait in a GWAS that includes both males and females present sexual dimorphism. To do this, we queried the LD-clumped genetic variants that were associated to each of our 530 traits at a $p < 1 \times 10^{-8}$ significance cut-off in a non-sex stratified effort using the UK Biobank data[4].

We found that the percentage of hits across the population (at $p < 1 \times 10^{-8}$) that present differences between the sexes (also at $p < 1 \times 10^{-8}$) ranges from 0 to 33.33% in binary traits and from 0 to 7.89% in non-binary traits. The average number of sdSNPs was 0.49% (SD 2.70%) in binary traits and 0.18% (SD 0.96%) in non-binary traits (Table S7). This indicates that sex dimorphism is not widespread across GWAS hits in sex-agnostic efforts.

If we look at it the other way around and question what percentage of sexually dimorphic hits are also significantly associated to a given trait in a non-sex stratified GWAS, we find that this ranges from 0 to 100% for both binary and non-binary traits. On average, this percentage is 67.46% (SD 45.23%) for binary traits and 23.10% (SD 29.59%) for non-binary traits (Table S7). This indicates that a fair portion of our dimorphic variants are also found to be significant in a sex-agnostic effort, but variability is quite big amongst traits.

#### SNP to gene analysis

##### *Traits with most genes*

A total of 93/103 phenotypes presented at least one dimorphic gene (combined $q_{2T} < 0.01$), with 76 and 78 phenotypes presenting at least one female and male dominant gene respectively (combined $q_F$ or $q_M < 0.01$). Traits with most dimorphic genes (combined $q_{2T} < 0.01$) are found in Table S11, the majority pertaining to the anthropometric class (including waist-hip circumference ratio, Figure S9, amongst others). Traits with most female or male dominant genes (Table S11) also showed a large predominance of anthropometric traits with the exception of several diseases like malabsorption/coeliac disease or hyperthyroidism showing large numbers of female dominant genes (87 and 83 respectively), and gout for male dominant genes (75). Interestingly, some traits presented only female or male dominant genes (13 and 17 traits respectively), including the aforementioned disease phenotypes, which are known to present differently in the sexes.

By visual inspection of our gene Manhattan plots (Figure S9 displays the one for Waist-hip circumference ratio), indications of LD having influenced our SNP to gene analysis are found, with nearby genes showing association to a given trait. This could influence our downstream analysis, potentially leading to enrichment of position-based gene sets, and as such is noted as a potential caveat.

##### *Genes with most traits*

A total of 3,556/18,253 genes were found to be dimorphic for at least one trait (combined $q_{2T} < 0.01$), with 1,863/18,253 and 1,402/18,253 genes being found to be female or male dominant for at least one trait respectively (combined $q_F$ or $q_M < 0.01$). Genes which were found to be dimorphic most recurrently are shown in Table S12. The 3 genes that showed up as dimorphic amongst most traits (considering $q_{2T} < 0.01$) were FRMD8 (chromosome 11), LOC101927789 (chromosome 11) and LSAMP (chromosome 3), with a total of 31, 29 and 29 traits respectively.

Notably, all top "female dominant" genes were found to be almost exclusively dominant in females across traits where they presented dimorphism, and vice-versa for the "male dominant" genes. This might suggest that male or female dominance is conserved across traits, i.e. a gene that presents dimorphism and has a larger effect in women will likely possess this larger effect in women across traits for which it was found to be dimorphic. It could also however be a result of the similarity of many phenotypes across UK Biobank, i.e. if the phenotype "Arm fat (left)" is affected dimorphically, "Arm fat (right)" will likely also show up as significant.

#### eQTL enrichment analysis

In order to assess whether the number of GxS interactions found was more than what you would expect than for non-sexually dimorphic variants, we repeated the analysis for these variants and their nearby genes (see Methods).

We then tested whether the number of GxS interactions found in our model was larger for sexually dimorphic SNPs than non-dimorphic SNPs by means of a Fisher's exact test, as explained in the Methods section. We focused on enrichment results for the $p < 1 \times 10^{-3}$ GxS significance threshold, as smaller thresholds provided a very small number of GxS interactions to perform the test (Table S19).

When considering a GxS significance threshold of $p < 1 \times 10^{-3}$, we found a total of 3 non-binary (artery coronary, liver, and skin not sun exposed suprapubic) and 4 binary traits (muscle skeletal, nerve tibial, cells transformed fibroblasts, and esophagus muscularis) presenting a Fisher's exact test *p*-value < 0.1. All of these tissues presented an odd's ratio (OR) > 2 for the presence of GxS in sexually dimorphic-gene pairs vs in non-sexually dimorphic gene pairs.

### Supplementary Figures

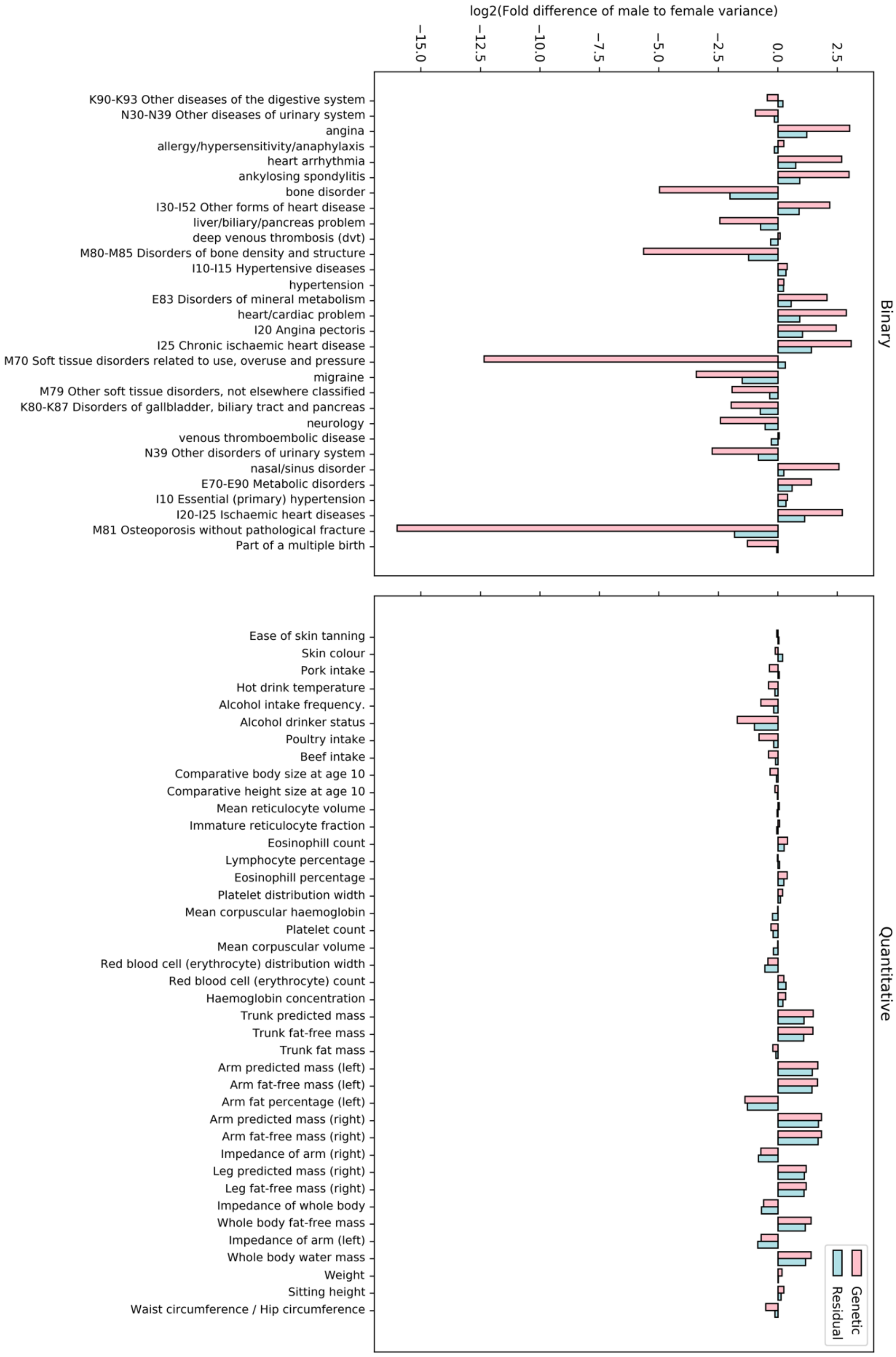

Figure S1. Barplot of variance fold difference between men and women for binary (top) and non-binary (bottom) traits with a significantly different heritability between the sexes at a $q < 0.05$ threshold. Pink bars represent fold change between the sexes in genetic variance, and blue bars represent fold change between the sexes in residual variance.

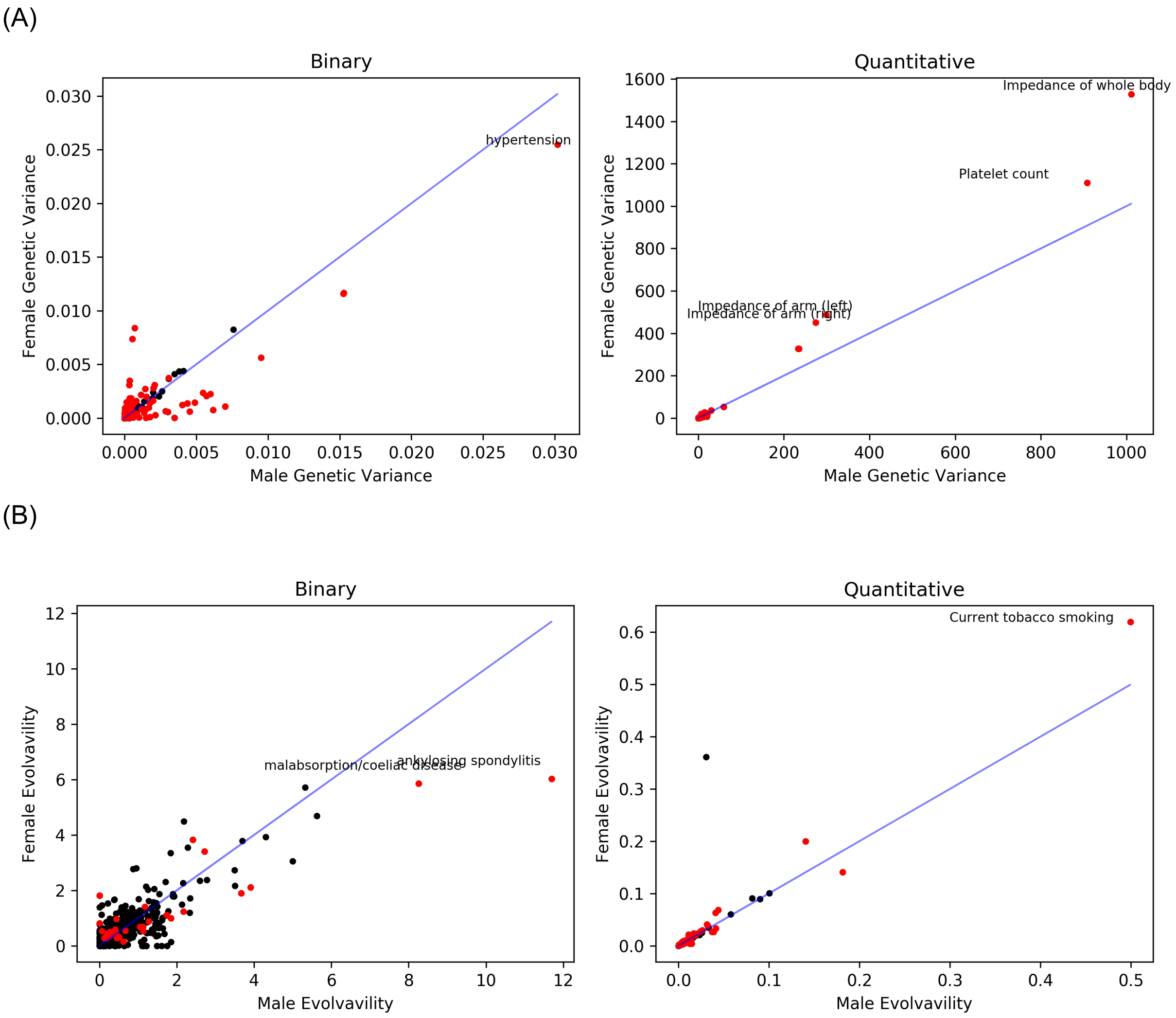

Figure S2. (A) Scatterplots comparing male genetic variance to female genetic variance for binary (left) and non-binary (right) traits. Each point represents a trait, and red points indicate traits for which the genetic variance between the sexes is significantly different ($q < 0.05$). (B) Scatterplots comparing male evolvability to female evolvability for binary (left) and non-binary (right) traits. Each point represents a trait, and red points indicate traits for which the evolvability between the sexes is significantly different ($q < 0.05$). Trait "basal metabolic rate" was removed as an outlier for both A and B.

(A)

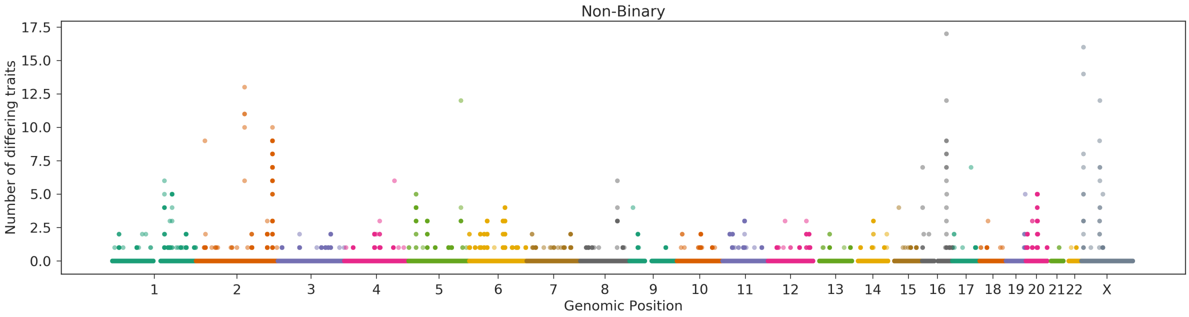

(B)

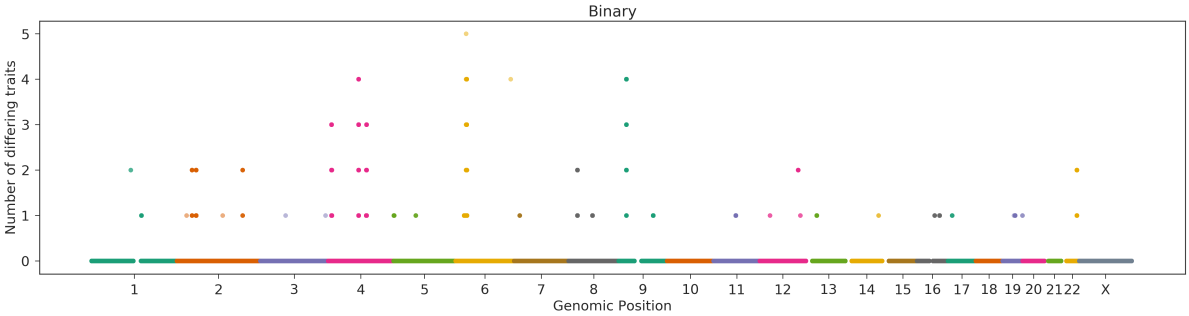

Figure S3. Manhattan plot of number of sdSNPs per genomic position in non-binary traits (each point represents a genetic variant, and its height the number of traits it affects in a sexually dimorphic manner) for (A) non-binary traits and (B) binary traits.

(A)

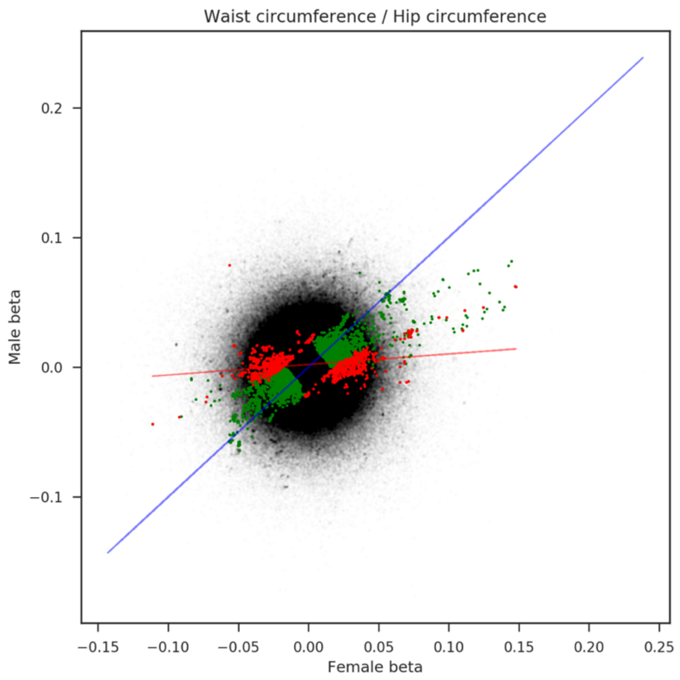

(B)

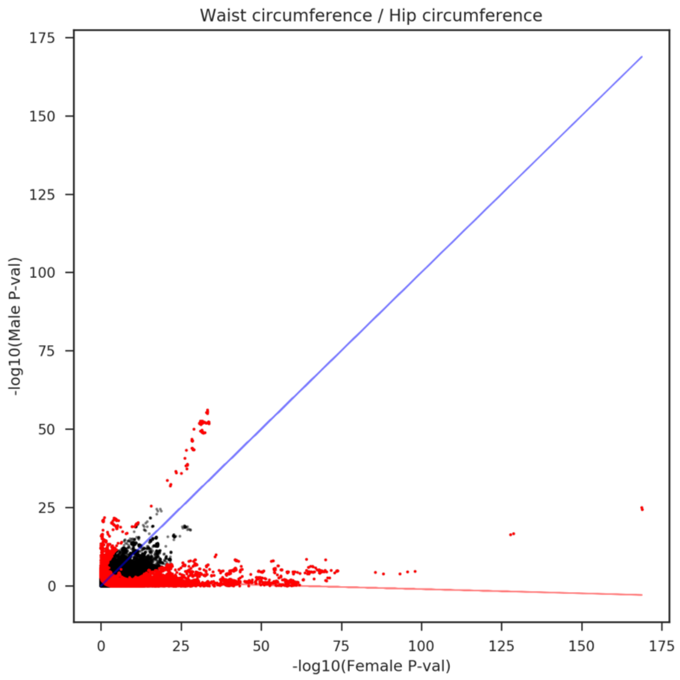

Figure S4. (A) Comparison of genetic effects in females versus males for waist-hip circumference ratio. Each point represents a genetic variant. In green are those found to be significant for the whole-population cohort (450K white-Europeans), and in red those that were found to be dimorphic at a $p < 1 \times 10^{-8}$ significance threshold. (B) Comparison of male and female *p*-values for significance of association for each variant across the autosomal genome to WHR. In red variants that were found to be dimorphic at a $p < 1 \times 10^{-8}$ significance threshold.

(A)

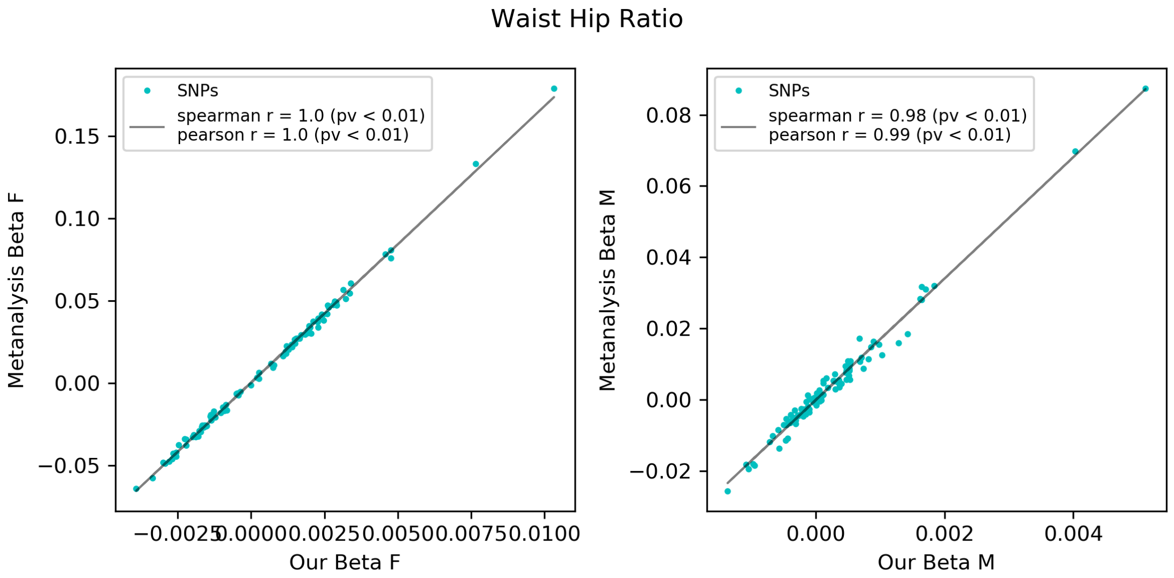
Waist Hip Ratio
SNPs
spearman r = 1.0 (pv < 0.01)
pearson r = 1.0 (pv < 0.01)
Metanalysis Beta F
Our Beta F
0.15
0.10
0.05
0.00
−0.05
−0.0025
0.0000
0.0025
0.0050
0.0075
0.0100
SNPs
spearman r = 0.98 (pv < 0.01)
pearson r = 0.99 (pv < 0.01)
Metanalysis Beta M
Our Beta M
0.08
0.06
0.04
0.02
0.00
−0.02
0.000
0.002
0.004

(B)

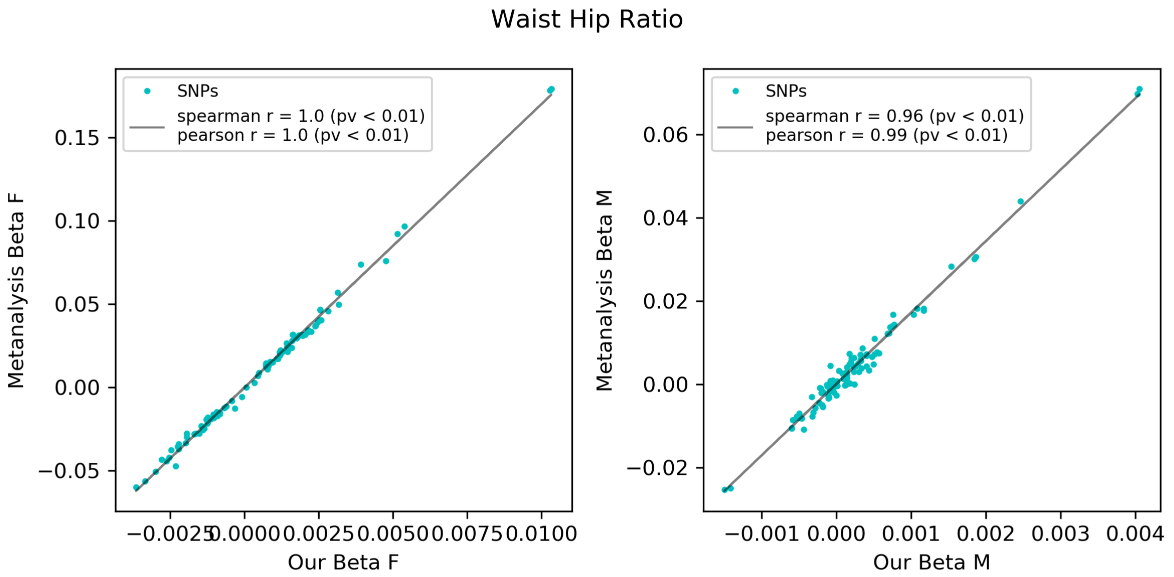
Waist Hip Ratio
SNPs
spearman r = 1.0 (pv < 0.01)
pearson r = 1.0 (pv < 0.01)
Metanalysis Beta F
Our Beta F
0.15
0.10
0.05
0.00
−0.05
−0.0025
0.0000
0.0025
0.0050
0.0075
0.0100
SNPs
spearman r = 0.96 (pv < 0.01)
pearson r = 0.99 (pv < 0.01)
Metanalysis Beta M
Our Beta M
0.06
0.04
0.02
0.00
−0.02
−0.001
0.000
0.001
0.002
0.003
0.004

(C)

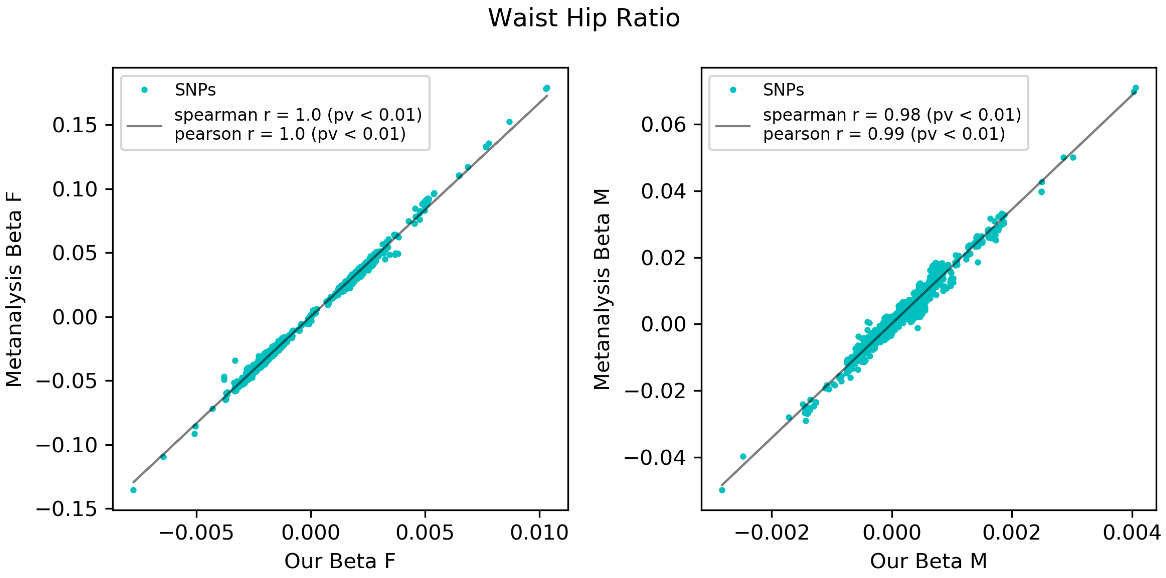

(D)

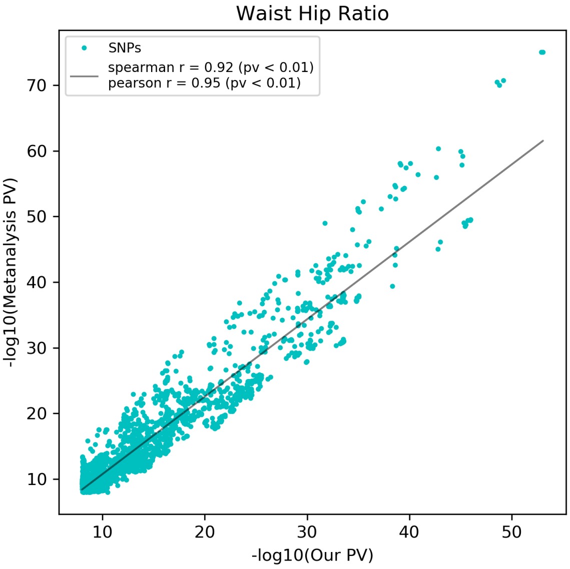

Figure S5. (A) Comparison of genetic effect estimates between our study and Pulit et al's, for our 100 lead sdSNPs in Waist-hip circumference ratio. Female estimates show on the left and male estimates on the right. (B) Comparison of genetic effect estimates between our study and Pulit et al's, for their 105 sdSNPs in Waist-hip circumference ratio. Female estimates show on the left and male estimates on the right. (C) Comparison of genetic effect estimates between our study and Pulit et al's, for their sdSNPs in Waist-hip circumference ratio, as calculated by us replicating our methodology using their summary statistics. Female estimates show on the left and male estimates on the right. (D) Comparison of our dimorphism test p-value to Pulit et al’s dimorphism test *p*-value, calculated by us using their summary statistics. SNPs considered are those that pass the $p < 1x10^{-8}$ significance cut-off for the dimorphism test using Pulit et al’s meta-analysis data.

(A)

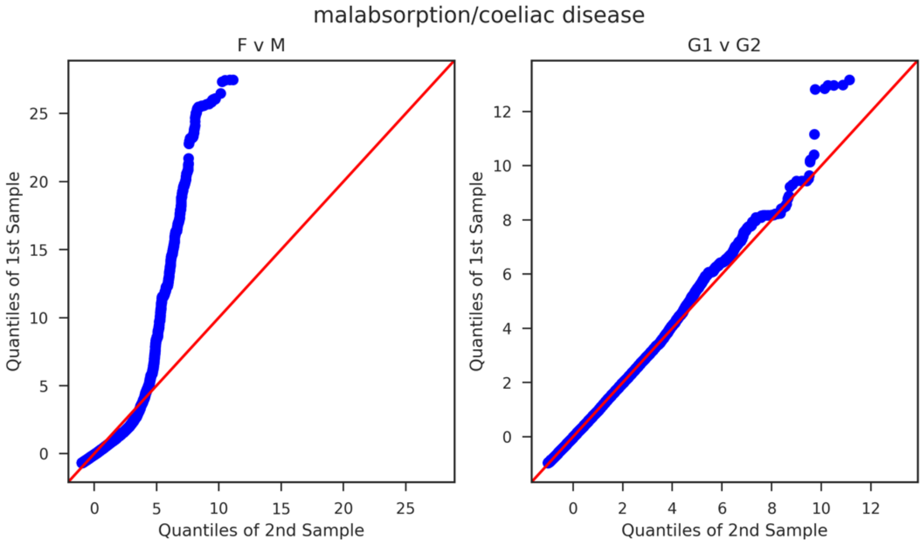

(B)

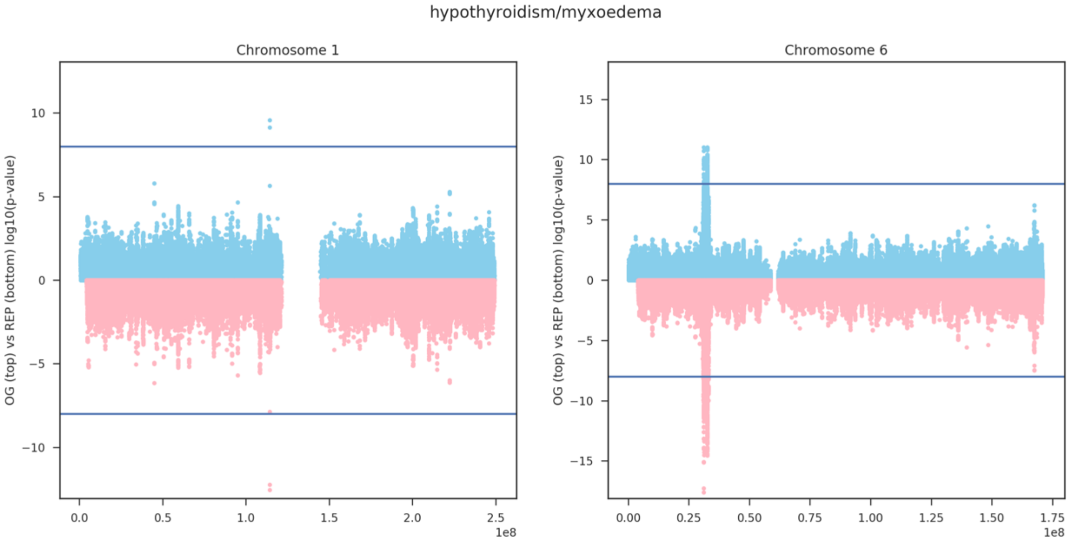

Figure S6. (A) QQ plots of our -log10 p-values corresponding to the comparison of genetic effects for the Female vs Male comparison on the left and the Group 1 vs Group 2 comparison on the right. (B) Mirrored Manhattan plot comparing sexual dimorphism log10 p-values for our original analysis (in blue) and the Female vs Male analysis (in pink), for hypothyroidism. Chromosome 1 is shown on the left, and Chromosome 6 on the right.

(A)

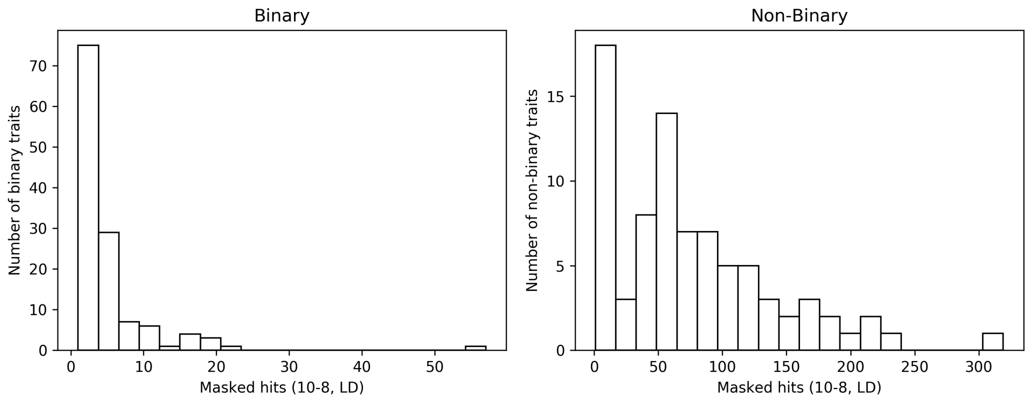
Binary
Number of binary traits
Masked hits (10-8, LD)
Non-Binary
Number of non-binary traits
Masked hits (10-8, LD)

(B)

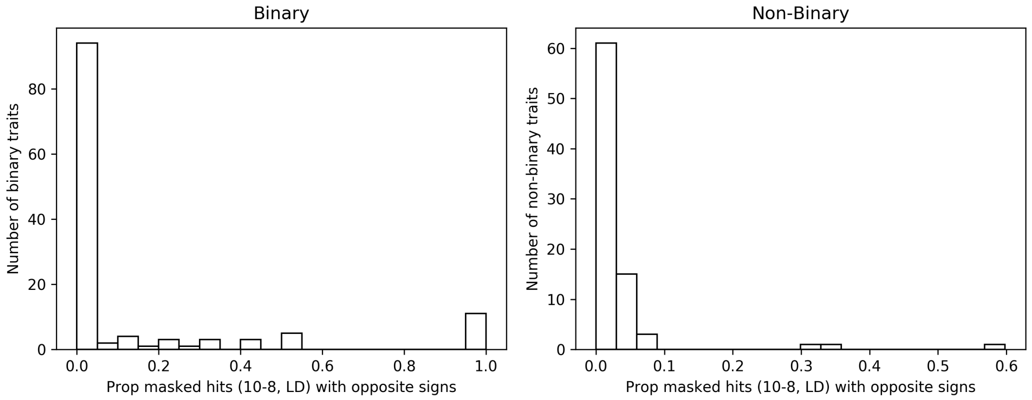
Binary
Number of binary traits
Prop masked hits (10-8, LD) with opposite signs
Non-Binary
Number of non-binary traits
Prop masked hits (10-8, LD) with opposite signs

(C)

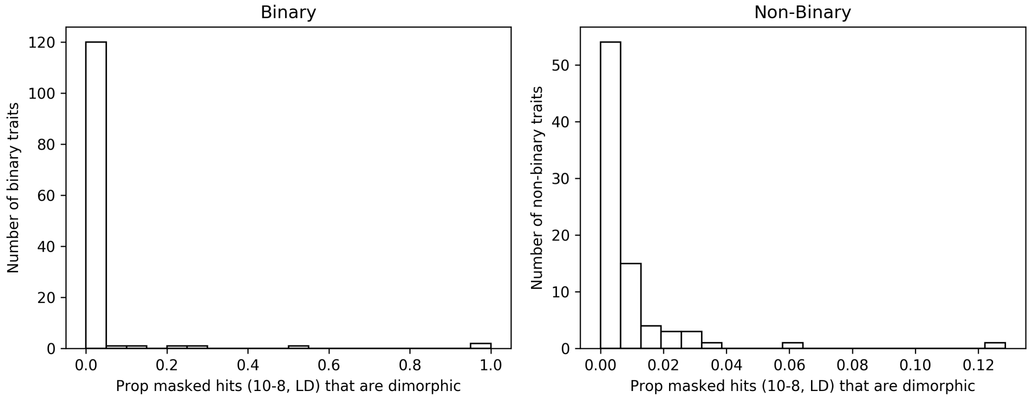
Binary
Number of binary traits
Prop masked hits (10-8, LD) that are dimorphic
Non-Binary
Number of non-binary traits
Prop masked hits (10-8, LD) that are dimorphic

(D)

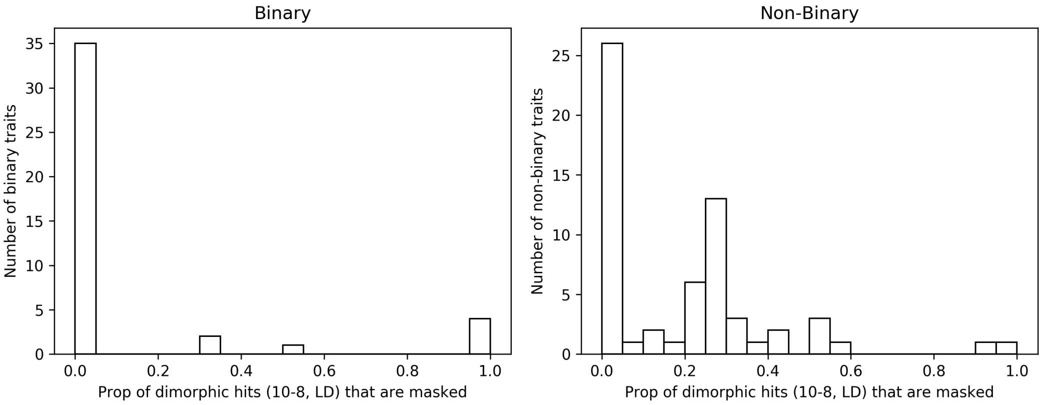

Figure S7. (A) Histogram of number of masked genetic variants found post LD clumping in binary (left) and non-binary (right) traits. (B) Histogram of proportion of masked SNPs (post-LD clumping) that presented opposite sign effects between the sexes. (C) Histogram of proportion of masked SNPs (LD clumped, $10^{-8}$ threshold) that were also found to possess significantly different genetic effects between the sexes for binary (left) and non-binary (right) traits. (D) Histogram of the proportion of sexually dimorphic SNPs that were found to be masked, for binary (left) and non-binary (right) traits.

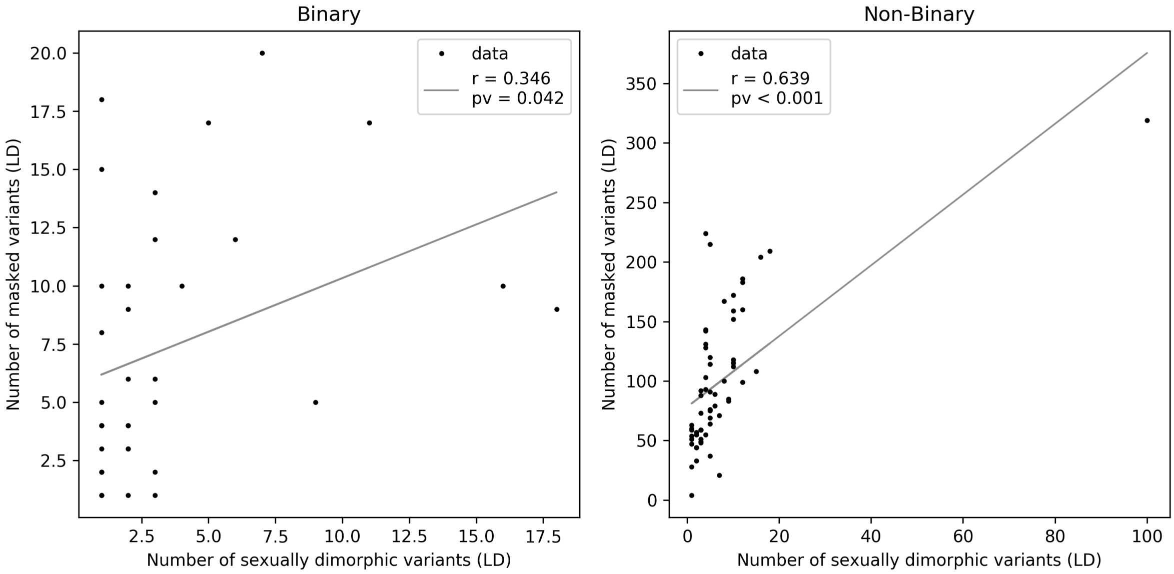

Fig S8. Scatterplot of number of sdSNPs versus number of masked variants ($p < 10^{-8}$, and LD clumped), for binary (left) and non-binary (right) traits.

(A)

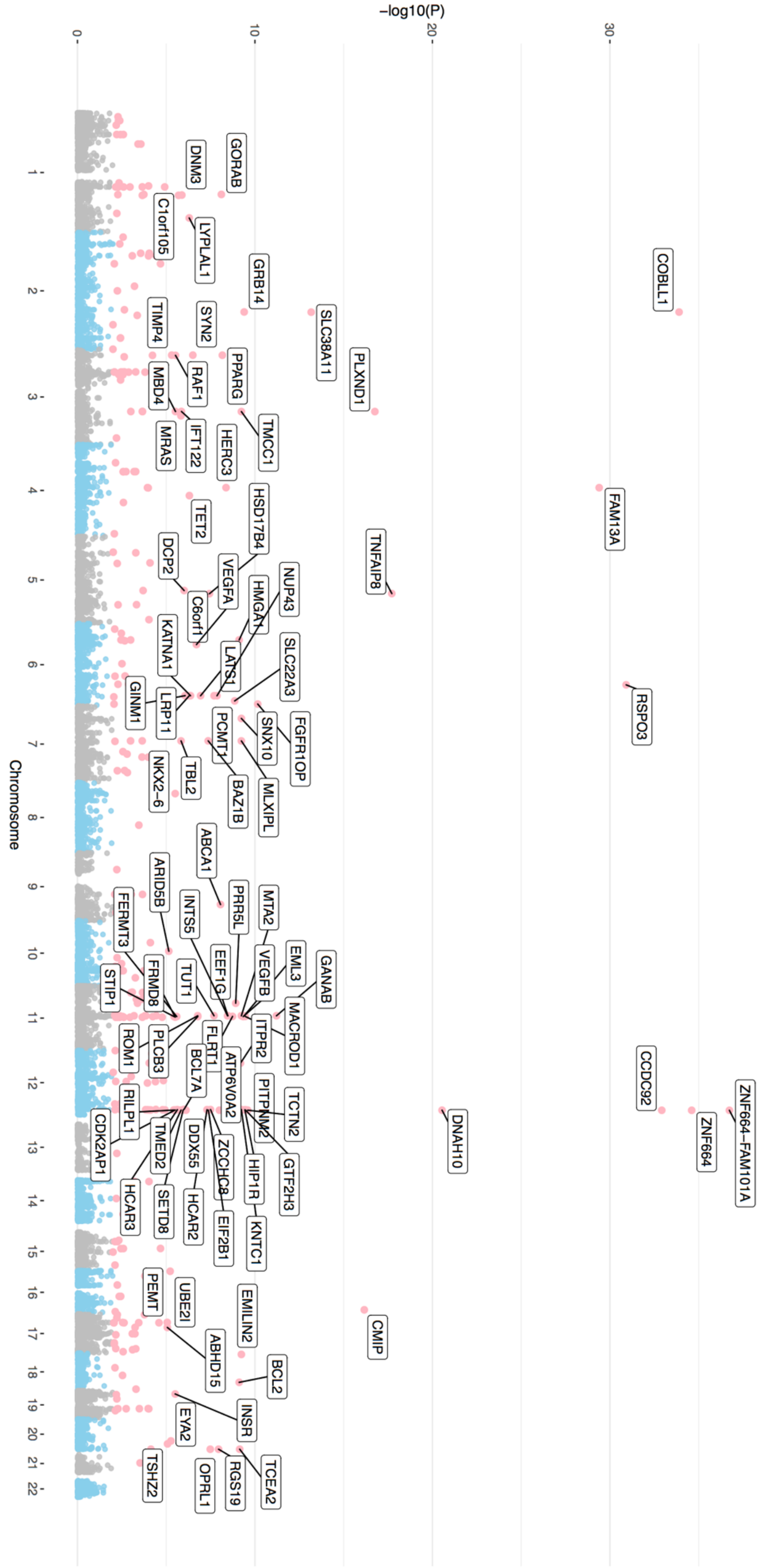

−log10(P)
0
10
20
30
Chromosome
GORAB
DNM3
C1orf105
LYPLAL1
GRB14
COBLL1
SLC38A11
SYN2
TIMP4
PPARG
RAF1
MBD4
PLXND1
IFT122
MRAS
HERC3
TMCC1
FAM13A
TET2
HSD17B4
DCP2
VEGFA
TNFAIP8
C6orf1
HMGA1
NUP43
KATNA1
LATS1
SLC22A3
GINM1
LRP11
RSPO3
PCMT1
SNX10
FGFR1OP
NKX2−6
TBL2
BAZ1B
MLXIPL
ABCA1
ARID5B
INTS5
PRR5L
MTA2
FERMT3
EEF1G
VEGFB
EML3
GANAB
STIP1
FRMD8
TUT1
ITPR2
MACROD1
ROM1
PLCB3
FLRT1
BCL7A
ATP6V0A2
PITPNM2
TCTN2
CCDC92
ZNF664−FAM101A
ZNF664
RILPL1
CDK2AP1
TMED2
DDX55
ZCCHC8
HIP1R
GTF2H3
DNAH10
HCAR3
SETD8
HCAR2
EIF2B1
KNTC1
PEMT
UBE2I
EMILIN2
CMIP
ABHD15
BCL2
EYA2
INSR
TSHZ2
OPRL1
RGS19
TCEA2

(B)

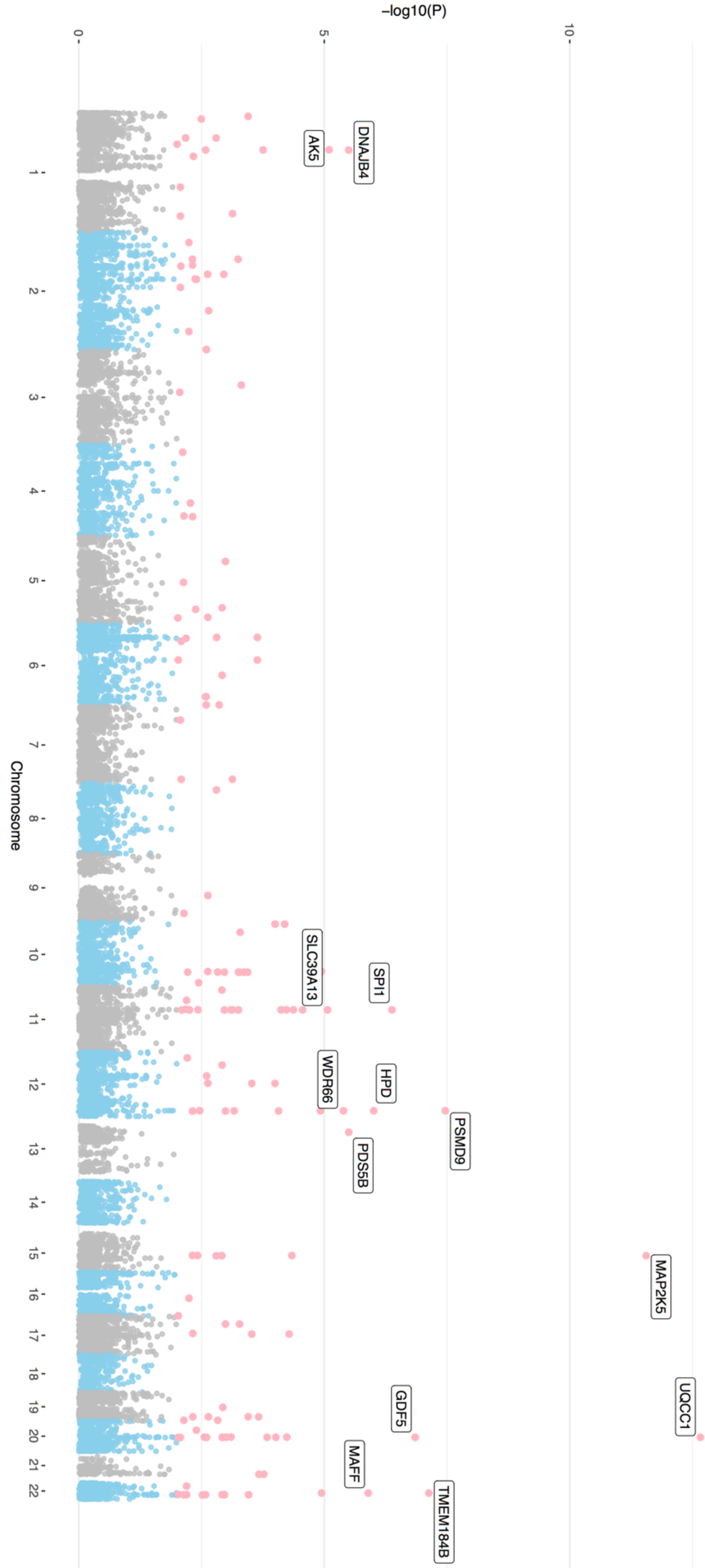

Figure S9. (A) Manhattan plot of Waist-hip ratio for gene-level analysis, using one-tail *p*-value for (A) females and (B) males.

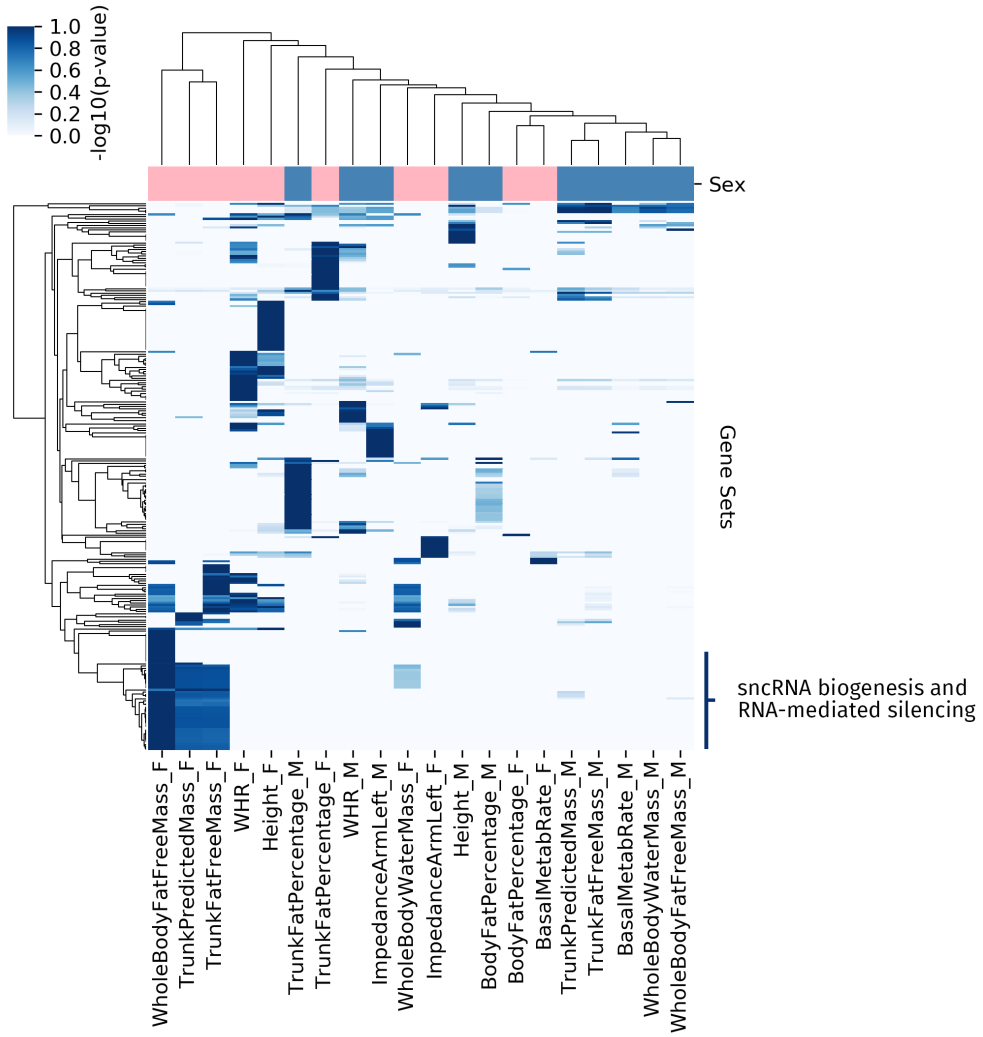

Figure S10. Heatmap with hierarchical clustering of FUMA set enrichment -log10 *p*-values for 251 gene sets that were found to be significantly differentially enriched between males and females (Fisher's exact test $q < 0.05$), as well as significantly differentially enriched to GWAS background genes (Fisher's exact test $q < 0.05$).

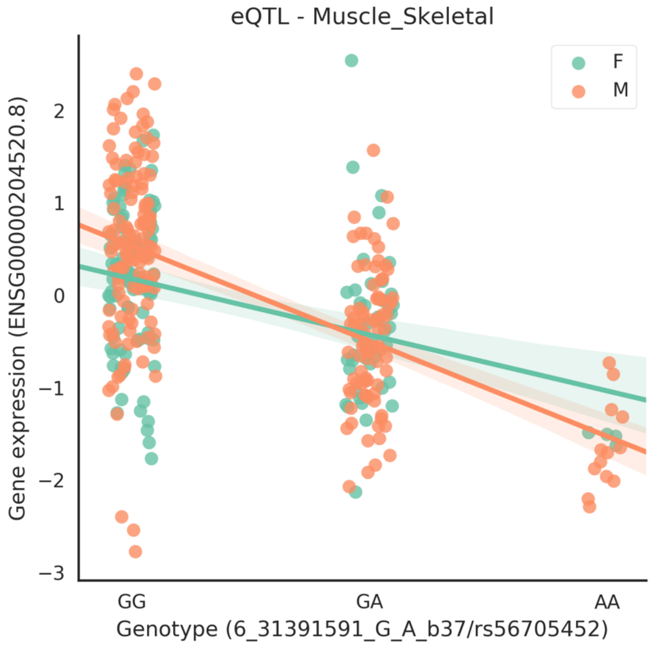

Figure S11. Relationship of genotype at variant rs56705452 with the expression of the transcript ENSG00000204520.8 in muscle skeletal tissue, for men (orange) and women (green).

### Supplementary Table Captions

Table S1. Full list of traits considered in study (530 in total), along with results for our heritability, genetic correlation and genetic effect comparison analyses for autosomal and X chromosome variants.

Table S2. Comparison of genetic correlation estimates for several traits with those that have been already published in the literature.

Table S3. Lead sdSNPs across traits.

Table S4. Comparison with GIANT for available traits, including comparison of sex-comparison p-values, female genetic effects and male genetic effects.

Table S5. Results of Model 1 and Model 2 replication.

Table S6. Results of randomization analysis for chromosome 1 and 6.

Table S7. Comparison to GeneATLAS results, including how many of our dimorphic SNPs are also significant in GeneATLAS and how many of the reported lead SNPs in GeneATLAS also present sexual dimorphism.

Table S8. Results of PGS comparison to phenotype values for 7 non-binary traits and 3 binary traits with over 10 dimorphic SNPs.

Table S9. Proportion of heritability explained by sdSNPs.

Table S10. Results of masking analysis across traits: including number of masked variants before and after LD clumping, number of these that present opposite sign genetic effects between the sexes, number of masked variants that are dimorphic, and number of dimorphic variants that are masked.

Table S11. Results of SNP to gene level results, with number of genes found to be significant considering $p_F$, $p_M$ and $p_{2T}$, per trait.

Table S12. Results of SNP to gene level results, with number of traits found to be significant considering $p_F$, $p_M$ and $p_{2T}$, per gene.

Table S13. Summary of results of FUMA (per trait) for our gene set enrichment analysis.

Table S14. Summary of results of FUMA (per gene set) for our gene set enrichment analysis.

Table S15. FUMA enrichment $p$-values for F, M and GWAS runs.

Table S16. eQTL results for autosomal dimorphic variant-gene pairs found in binary traits.

Table S17. eQTL results for autosomal dimorphic variant-gene pairs found in non-binary traits.

Table S18. eQTL results for X-chromosome dimorphic variant-gene pairs found in non-binary traits.

Table S19. eQTL enrichment for non-binary and non-binary traits.
